# macroH2A2 shapes chromatin accessibility at enhancer elements in glioblastoma to modulate a targetable self-renewal epigenetic network

**DOI:** 10.1101/2021.02.23.432465

**Authors:** Ana Nikolic, Anna Bobyn, Francesca Maule, Katrina Ellestad, Xueqing Lun, Michael Johnston, Christopher J Gafuik, Franz J Zemp, Seungil Paik, Nicoletta Ninkovic, Sajid A Marhon, Parinaz Mehdipour, Yaoqing Shen, N. Daniel Berger, Duncan K Brownsey, Peter B Dirks, Darren J Derksen, Steven JM Jones, Daniel de Carvalho, Donna L Senger, Jennifer A Chan, Douglas J Mahoney, Marco Gallo

## Abstract

Self-renewal is a crucial property of glioblastoma cells and is enabled by the choreographed function of chromatin regulators and transcription factors. Identifying targetable epigenetic mechanisms of self-renewal could represent an important step toward developing new and effective treatments for this universally lethal cancer. Here we uncover a targetable epigenetic axis of self-renewal mediated by the histone variant macroH2A2. Using patient-derived *in vitro* and *in vivo* models, we show that macroH2A2 has a direct role in shaping chromatin accessibility at enhancer elements to antagonize transcriptional programs of self-renewal. Pharmaceutical inhibition of the chromatin remodeler Menin increased macroH2A2 levels and repressed self-renewal. Our results reveal a targetable epigenetic mechanism of self-renewal controlled by macroH2A2 and suggest new treatment approaches for glioblastoma patients.

**SIGNIFICANCE:** Glioblastoma is an incurable brain cancer. Malignant self-renewing cells have been shown to drive tumor growth, to be refractory to current treatment approaches and to seed relapses, which ultimately prove lethal. Identifying new and targetable mechanisms associated with self-renewal could be a fundamental first step in designing effective therapies that slow or prevent glioblastoma relapses. Using patient-derived models of glioblastoma, we deployed epigenomic approaches and functional assays to define the role of the histone variant macroH2A2 in repressing self-renewal. We identified compounds that increase macroH2A2 levels and repress self-renewal, including a Menin inhibitor. As Menin inhibitors are being tested in clinical trials, these compounds could be used in new therapeutic paradigms to target self-renewing cell populations in glioblastoma.

## INTRODUCTION

Glioblastoma (GBM) is the most common malignant primary brain tumor in adults and carries abysmal prognoses, even in cases treated with gross total resection and adjuvant chemoradiotherapy (Stupp et al., 2005). Several factors are thought to contribute to the aggressiveness of GBM, especially at recurrence: The activation of drug resistance mechanisms (Chen et al., 2012), hypermutation (Cahill et al., 2007; Hunter et al., 2006; Yip et al., 2009), a highly immunosuppressive tumor microenvironment (Alvarado et al., 2017; Antonios et al., 2017), and underlying disease characteristics, most notably a high degree of intratumoral heterogeneity (Neftel et al., 2019; Suvà et al., 2014). Recent studies have shown that the cellular heterogeneity observed in GBM occurs both at the genetic and epigenetic level (Gallo et al., 2015; Lan et al., 2017; Neftel et al., 2019; Suvà et al., 2014). It is thought that epigenetic heterogeneity in GBM reflects the co-existence of functionally non-equivalent populations of cells, similarly to what has been observed in normal developmental paradigms. This functional interpretation of epigenetic heterogeneity in GBM well aligns with the experimentally-validated concept that GBM is populated by cell populations with different functional properties, including slow-cycling self-renewing cells, proliferative progenitor-like cells with limited self-renewal and differentiated progeny that are destined to die (Cusulin et al., 2015a; Lan et al., 2017; Lathia et al., 2010; Miller et al., 2017; Singh et al., 2004; Suvà et al., 2014) .

GBM self-renewing cells (GSCs) are expandable in culture (Pollard et al., 2009), can differentiate into non-self-renewing cell types *in vitro* (Galli et al., 2004; Singh et al., 2003) and are capable of tumor initiation and serial propagation of the tumor in immunocompromised mice *in vivo* (Singh et al., 2004). Studies have shown GSCs are more resistant to the standard of care treatment for GBM than non-self-renewing cells, and they represent a larger fraction of tumor cells at relapse than at diagnosis (Bao et al., 2006; Liu et al., 2006). GSCs therefore play crucial roles in tumor growth, therapy resistance and in seeding relapses. However, therapeutic targeting of self-renewing cells is currently an unmet clinical need.

Pro-self-renewal epigenetic programs are maintained by the choreographed function of several chromatin remodelers. These include the histone methyltransferase DOT1L and arginine demethylase JMJD6 (MacLeod et al., 2019), members of the mixed-lineage leukemia (MLL) family (Gallo et al., 2013; Heddleston et al., 2012) and of the polycomb-repressive complexes 1 and 2 (PRC1/2) (Abdouh et al., 2009; Suvà et al., 2009), among others. These chromatin remodelers promote self-renewal by activating transcriptional networks associated with stemness and by repressing differentiation. This can be achieved through the activation of master transcription factors (Rheinbay et al., 2013; Suvà et al., 2014) or by modulating the expression of histone variants, which ultimately shape chromatin organization and downstream transcriptional programs .

Histone variants differ from core histones at key amino acids that determine their patterns of incorporation into chromatin and their effects on transcription of downstream genes (Maze et al., 2014). For instance, *H2AFY* and *H2AFY2* encode two closely related variants of core histone H2A, namely macroH2A1 and macroH2A2, respectively. They share only about 60% sequence identity in their histone domain with core histone H2A (Buschbeck and Hake, 2017), and contain two additional domains: A basic linker region with putative DNA binding function, and a “macro” domain with little sequence conservation between the two paralogs (Costanzi and Pehrson, 2001). Because of the large size of the macro domain, substitution of H2A with macroH2A1 or macroH2A2 is predicted to have large effects on chromatin organization (Cantariño et al., 2013). Loss or downregulation of macroH2A1 and macroH2A2 has been described in a number of malignancies, including bladder cancer, melanoma, lung cancer, and gastric cancer (Kapoor et al., 2010; Novikov et al., 2011; Sporn et al., 2009).

MacroH2A1 was first studied for its role in X chromosome inactivation in female mammalian cells (Costanzi and Pehrson, 1998). It localizes to large chromatin domains throughout the genome that encompass developmentally regulated genes and regions of imprinting, and it accumulates at senescence-associated heterochromatic foci (Chen et al., 2015; Choo et al., 2006; Gamble et al., 2010; Zhang et al., 2005, 2007). Both macroH2A1 and macroH2A2 are involved in the repair of double-strand breaks, and knockdown of macroH2A1 impairs DNA double strand repair (Kim et al., 2017, 2018; Kozlowski et al., 2018; Timinszky et al., 2009). MacroH2A2 loss has been described in invasive melanoma (Kapoor et al., 2010). Knockdown or knockout of macroH2A paralogs increase reprogramming efficiency of somatic cells into induced pluripotent stem cells (Barrero et al., 2013), suggesting that these histone variants might be involved in repression of stemness properties, including self-renewal.

The molecular mechanisms exploited by macroH2A paralogs in modulating stemness and self-renewal are not currently understood, especially in the context of cancer. Given the current inability to target self-renewing cells in most cancer types, dissection of these mechanisms could be important to identify new treatment options for GBM and other malignancies characterized by intratumoral functional heterogeneity. Here we deploy epigenomic and functional assays to unravel the mechanisms utilized by macroH2A2 to control self-renewal programs in GBM, and pursue potential translational avenues with our preclinical models.

## RESULTS

### Low expression of *H2AFY2* is a negative prognostic factor for GBM patients

We embarked on a systematic analysis to identify genes encoding histone variants that may be involved in GBM biology. Our approach consisted in looking for histone variant genes whose expression could stratify GBM patients based on overall survival. One of our best candidates was *H2AFY2*, the gene encoding the histone H2A variant macroH2A2. Median expression levels were used to classify patients as high- or low-expressors. We found that low levels of *H2AFY2* transcription were associated with shorter overall survival in IDH-wildtype GBM in the Gravendeel cohort (Gravendeel et al., 2009) (log-rank p = 0.0069), as well as in the GBM cohort collected by The Cancer Genome Atlas (TCGA, 2015) (**Figure 1A, 1B**). Age is a potent confounder for overall survival, with younger glioma patients usually performing better than older patients. A multivariate Cox regression model also showed that increased *H2AFY2* expression had prognostic significance in both cohorts (hazard ratio: 0.54 [0.38 – 0.78]; **Figure 1C; Figure S2A-S2C**), even when adjusted for known prognostic factors such as IDH mutation status, age, and treatment status **(Figure S2C)**. In the TCGA cohort, the survival benefit associated with high *H2AFY2* transcription was also observed in patients with recurrent disease, and these patients were noted to have significantly higher levels of *H2AFY2* expression on average (**Figure S2D; Figure S2E).** Transcription levels of *H2AFY* (which encodes macroH2A1) had no prognostic significance in either patient cohort **(Figure S1B, S1C).** Moreover, the effects of *H2AFY2* appeared to be specific to IDH-wildtype GBM, as no effect was seen in IDH-mutant tumours in the Gravendeel dataset **(Figure S1D).** When RNA-seq data from the TCGA are separated by transcriptional subtype, the survival benefit of *H2AFY2* expression is significant specifically in proneural tumours (p = 0.0024; log-rank test) and not in classical or mesenchymal tumours (**Figure 1D)**.

**Figure 1.**
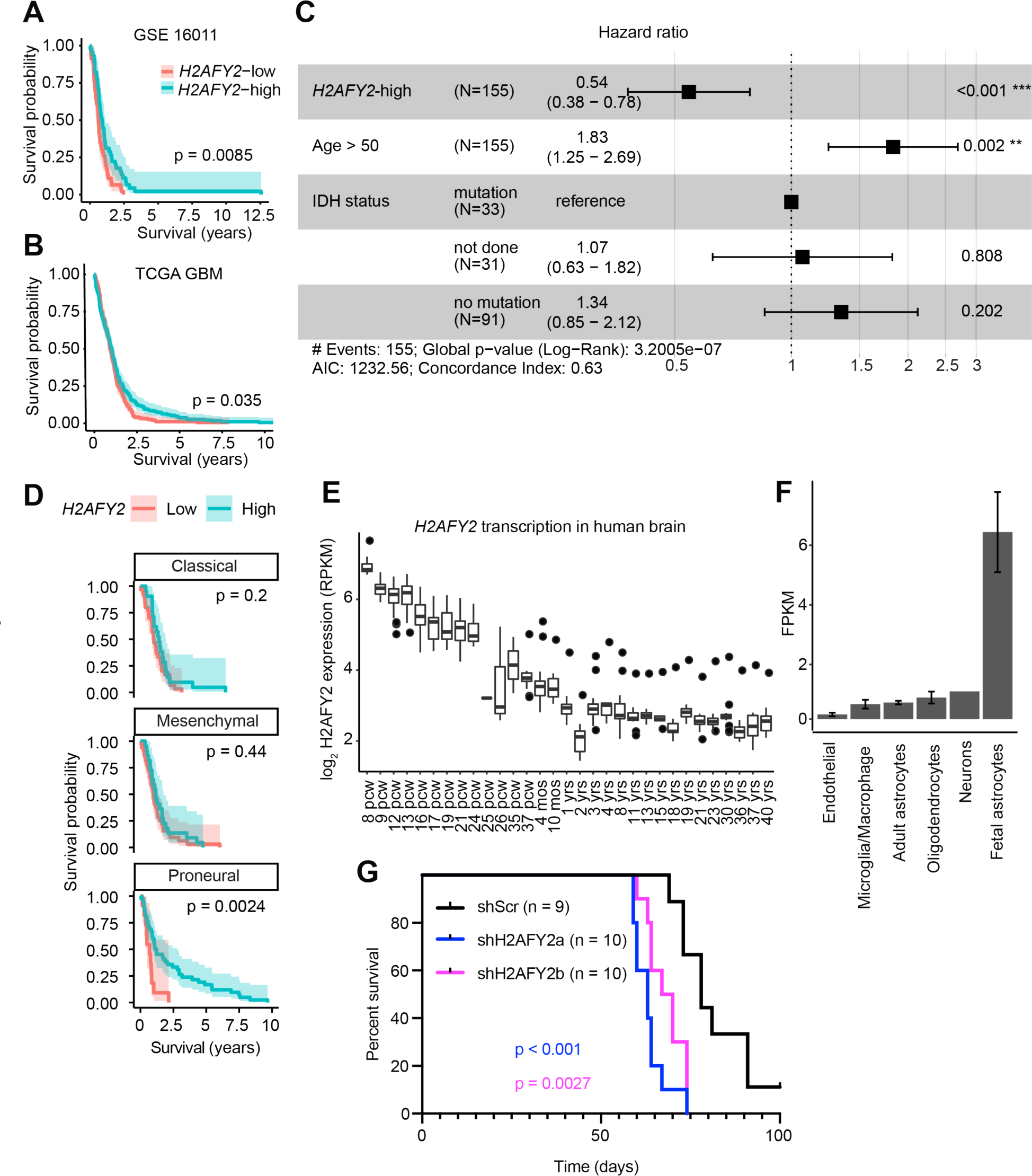
Low transcription levels of *H2AFY2*/macroH2A2 contribute to GBM aggressiveness. **(A, B)** Kaplan-Meier survival analysis of adult *IDH*-wildtype glioblastoma patients (GSE16011; Gravendeel et al 2009; TCGA glioblastoma cohort) based on *H2AFY2* transcription levels. *H2AFY2-*low and -high groups were determined by median gene expression. Shaded region represents 95% confidence interval. P value was obtained by log-rank test. **(C)** Hazard ratios for macroH2A2 expression in a multivariate Cox regression model adjusting for other factors relevant for glioblastoma (age, IDH mutation status). Error bars represent 95% confidence intervals. **(D)** Kaplan-Meier survival analysis of adult glioblastoma patients separated by transcriptional subtype (TCGA). Shaded region represents 95% confidence interval. P values obtained by log-rank test.**E)** Expression of *H2AFY2* at different timepoints of brain development. Datasets were accessed through the BrainSpan database. **(F)** Transcriptional levels of *H2AFY2* were assessed in different human brain cell types. Datasets were accessed through the BrainSeq2 database. **G** Orthotopic xenograft experiments to assess the effects of *H2AFY2* knockdown on survival of transplanted mice. Patient-derived GSCs carrying either scrambled control shRNA constructs (shScr; n = 9) or independent shRNAs targeting *H2AFY2* (shH2AFY2a/b; n = 10 mice per group) were transplanted orthotopically in immunocompromised mice. P values were calculated with the log-rank test.

When expression of *H2AFY2* is compared to expression of *H2AFY* in the TCGA glioblastoma cohort, the former has a clear bimodal distribution, a pattern not seen for *H2AFY* (macroH2A1) **(Figure S1E)**. These data point to a specific role for macroH2A2 in GBM chromatin biology.

We used transcriptome datasets from the Allen Brain Atlas BrainSpan database (Miller et al., 2014) to extract information on *H2AFY2* expression during human brain development in samples collected between 8 weeks post-conception and 40 years of age. We found that *H2AFY2* expression peaks at 8 weeks post-conception, steadily declines during fetal development and stabilizes at 2-3 years of age into adulthood (**Figure 1E**). Our analyses of data from BrainSeq2 also show that expression of *H2AFY2* is particularly elevated in fetal astrocytes compared to other brain cell types (**Figure 1F**) (Zhang et al., 2016). Single-cell expression data from mouse striatum highlights a fetal expression pattern for *H2afy2*, with preferential expression largely in neuronal precursor cells and oligodendrocyte precursor cells **(Figure S1C)** (Saunders et al., 2018). Transcription of *H2AFY2* in the brain therefore appears to be temporally-regulated, with the highest expression observed during early fetal development in both humans and mice.

Because *H2AFY2* levels have prognostic significance and are dynamically regulated during brain development, we decided to investigate whether this gene plays a direct role in the biology of GBM. We used our patient-derived GSC cultures to generate stable inducible *H2AFY2* knockdown models of GBM. We transplanted GSCs carrying control scrambled shRNA constructs (shScr) or either of two shRNAs targeting *H2AFY2* (shH2AFY2a/b) into the forebrains of NSG mice. We observed reproducible negative effects of *H2AFY2* knockdown on mouse survival (log-rank p < 0.001 for shH2AFY2a and p = 0.0027 for shH2AFY2b) (**Figure 1G**). Therefore our patient-derived knockdown models recapitulate the association between low *H2AFY2* levels and poor prognosis we observed in glioma patient cohorts (**Figures 1A,1B**). Importantly, our results indicate that this gene plays a direct functional role in GBM biology and deserves further investigation.

### macroH2A2 antagonizes self-renewal in GBM

We previously reported that the histone H3 variant H3.3 has a strong role in repressing self-renewal in GBM (Gallo et al., 2015). Self-renewal, or self-replication, is a key property of a population of cells that propagate tumor growth (Magee et al., 2012). It was shown that the fraction of GBM cells that can self-renew, as well as transcriptional signatures associated with self-renewal, are negative prognostic factors for this brain cancer (Murat et al., 2008). Given the significance of self-renewal in GBM, we investigated whether macroH2A2 plays a role in modulating this crucial functional property. The frequency of self-renewing cells in a population can be estimated with *in vitro* limiting dilution assays (LDAs), which use sphere-forming frequency as a measure of self-renewal. We used our doxycycline-inducible *H2AFY2* knockdown models (**Figure 2A,B; Figure S2A**) for *in vitro* LDAs in three different primary patient-derived cultures (G523, GSC3,GSC5), and found that knocking down *H2AFY2* resulted in increased self-renewal (**Figure 2C-2E**). These *H2AFY2* knockdown constructs were specific to this H2A variant and showed no effect on protein levels of the related variant macroH2A1 **(Figure S2B)**. In order to cross-check these results with an independent experimental system, we decided to overexpress *H2AFY2* using a fusion protein composed of catalytically dead Cas9 (dCas9) and the transcriptional activator VPR (dCas9-VPR system (Chavez et al., 2015); **Figure 2F**). We transduced GSCs with a lentivirus carrying dCas9- VPR to generate a stable inducible overexpression model. We then transfected this GSC line with a pool of five single guide RNAs (sgRNAs) targeting *H2AFY2*. RT-qPCR showed that this experimental system resulted in a 10-fold overexpression of *H2AFY2* over control cells transfected with scrambled sgRNAs (sgScr), with no effects on transcription levels of the close paralog *H2AFY* encoding the H2A variant macroH2A1 (**Figure 2G; Figure S2C, S2D**). We therefore employed this *H2AFY2*-specific overexpression models in *in vitro* LDAs, which showed that *H2AFY2* overexpression causes a ∼50% reduction in sphere-forming frequency (**Figure 2H**). Therefore, our knockdown and overexpression systems concordantly show an antagonistic effect of macroH2A2 on self-renewal in patient-derived GSC cultures.

**Figure 2.**
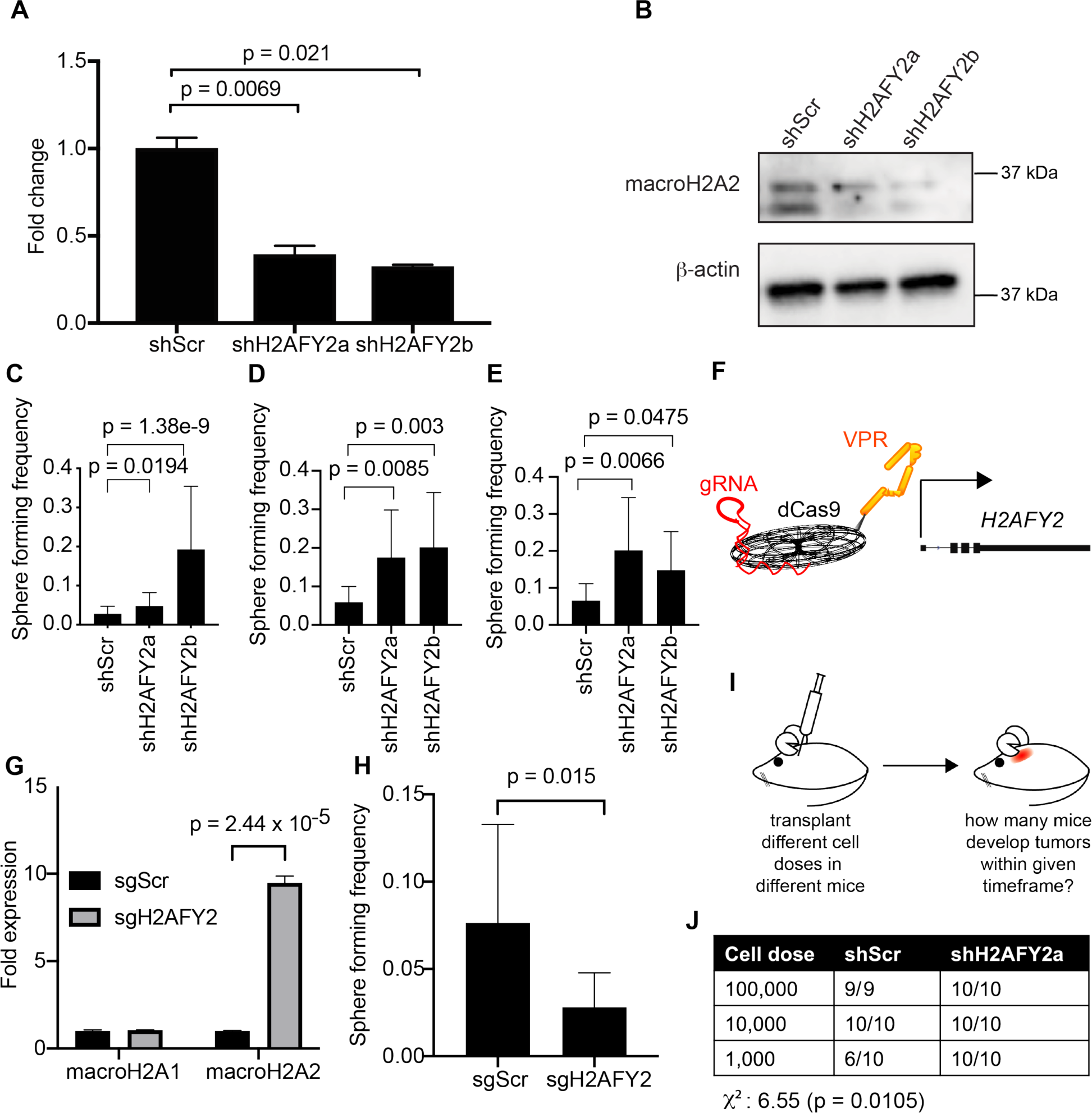
macroH2A2 antagonizes self-renewal in GBM. **(A)** qRT-PCR to assess the transcriptional levels of *H2AFY2* in cells carrying control (shScr) or knockdown (shH2AFY2a/b) constructs 48 hours after doxycycline induction. Expression normalized to actin and GAPDH. P value were determined with a two-tailed t-test. The experiment was repeated 4 times. **(B)** Western blot to compare macroH2A2 protein levels in control and knockdown cells after 14 days of shRNA induction. The experiment was repeated 3 times. **(C-E)** Limiting dilution assay results after 14 days of doxycycline induction in G523 glioma cells **(C)**, GSC3 **(D)**, and GSC5 **(E)**. P value was determined by Chi-square test with the tool ELDA (see Methods). Error bars: 95% confidence interval. Statistics from 6 technical replicates; the experiment was repeated 3 times. **(F)** Schematic of the dCas9-VPR overexpression system. **(G)** Validation of overexpression system by qRT-PCR after 48 hours of induction by sgRNA. Transcription was normalized to actin and GAPDH. P value was determined with a two-tailed t-test. The experiment was repeated 2 times. **(H)** Limiting dilution assay of G523 cells with dCas9-VPR and guide RNA targeting macroH2A2 or a non-targeting guide control at 7 days. P value was determined by Chi-square test with the tool ELDA (see Methods). Error bars: 95% confidence interval. Statistics from 6 technical replicates; the experiment was repeated 3 times. **(I)** Schematic of *in vivo* limiting dilution assay. **(J)** Overview of *in vivo* limiting dilution assay results. Mice were transplanted orthotopically with either shScr or shH2AFY2a-transduced GSCs. P value and chi square value obtained by Chi-square test.

To further validate these findings, we performed *in vivo* LDAs. We transplanted GBM cells carrying our doxycycline-inducible stable *H2AFY2* system or shScr controls into the forebrains of NSG mice (**Figure 2I**). We transplanted 10 mice at each cell dose (100,000, 10,000 and 1,000 cells) for knockdown and control cells. At the lowest dose, 10 out of 10 mice transplanted with *H2AFY2*-knockdown cells developed tumors, whereas only 6 out of 10 control mice did (**Figure 2J**). Overall, we observed a significant difference in engraftment potential between GBM cells with *H2AFY2* knockdown and control cells (χ^2^ p = 0.0105). Our *in vivo* LDAs therefore confirm our *in vitro* experiments and support an antagonistic role of macroH2A2 on self-renewal.

### macroH2A2 represses a transcriptional signature of self-renewal in GBM

Given that our *in vitro* and *in vivo* functional assays showcased a role for macroH2A2 in repressing self-renewal properties of GBM cells, we hypothesized that this histone variant might regulate expression of stemness genes. To test this hypothesis, we performed RNA-seq on GBM cells with *H2AFY2* knockdown (two independent hairpins) or shScr constructs (two biological replicates per condition). We then looked at the effects of *H2AFY2* knockdown on a recently published GBM stemness signature composed of 17 genes (Suvà et al., 2014). We found that overall, *H2AFY2* knockdown resulted in increased transcription of these stemness genes, with strong and consistent effects especially on *VAX2*, *SALL2*, *SOX8*, *SOX21*, *POU3F*3, *PDGFRA*, *ASCL1* and *OLIG2* (**Figure 3A**). These data support a role for macroH2A2 in repressing self-renewal by antagonizing a stemness transcriptional network in GBM.

**Figure 3.**
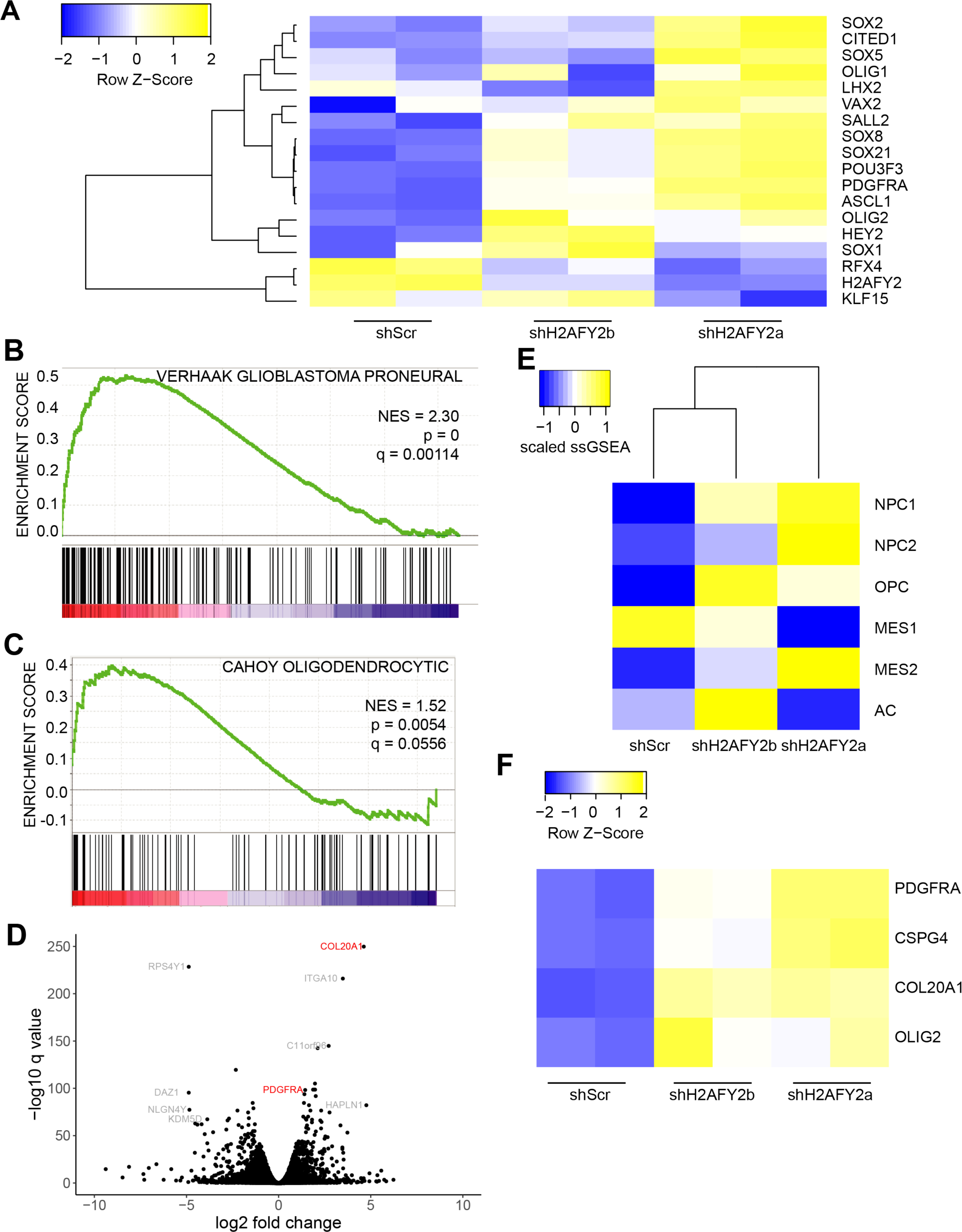
MacroH2A2 knockdown leads to widespread gene expression changes in stemness and oligodendroglial genes. **(A)** RNA-seq was used to determine transcriptional levels of an established stemness gene signature at 7 days following *H2AFY2*/macroH2A2 knockdown.Two biological replicates were used per condition. **(B,C)** Two signatures enriched by GSEA in knockdown versus control cells. **(D)** Volcano plot highlighting differentially expressed genes after 7 days of *H2AFY2*/macroH2A2 knockdown in G523 cells. **(E)** Comparative ssGSEA analysis of the Neftel scRNA-seq GBM transcriptional subtypes (Neftel et al., 2019) in control versus *H2AFY2* knockdown samples. **(F)** Heatmap showing differential expression of oligodendrocyte precursor cell-associated genes in knockdown cells versus control. Two biological replicates per condition.

Gene set enrichment analysis (GSEA) (Subramanian et al., 2005) of differentially expressed genes upon *H2AFY2* knockdown revealed positive enrichment for signatures associated with the proneural molecular subtype of GBM (Verhaak et al., 2010) (**Figure 3B**) and with oligodendrocytic gene signatures (**Figure 3C**). Volcano plots further illustrated that some of the most upregulated genes upon *H2AFY2* knockdown are oligodendrocyte markers, including *COL20A1* and *PDGFRA* (**Figure 3D**) (Rivers et al., 2008). When gene expression is compared to the single-cell transcriptional subtypes identified by Neftel et al, we see an increase in the OPC and NPC1 lineages by ssGSEA, and a reduction in mesenchymal (MES1) and astrocytic (AC) lineages (**Figure 3E**). Interestingly, with shH2AFY2a, which has a stronger knockdown, we see more of the MES and NPC2 phenotypes, which are not seen with the shH2AFY2b knockdown. This suggests that macroH2A2 may act as a rheostat regulating the transitions between distinct cellular fates. We also found that *CSPG4/NG2*, a widely-used marker of the oligodendrocyte lineage, is transcribed at higher levels following *H2AFY2* knockdown (**Figure 3F**). Overall, our data point to a role for macroH2A2 in repressing gene expression signatures associated with self-renewal and the oligodendrocytic lineage. This is a particularly intriguing finding because oligodendrocyte progenitor cells have been proposed as the cell of origin of GBM based on experimental work with murine (Alcantara Llaguno et al., 2015; Liu et al., 2011) and patient-derived (Yuan et al., 2018) models.

### MacroH2A2 maintains chromatin organization at developmentally regulated genes

MacroH2A variants have been mostly studied in the context of their roles in chromatin compaction, including X inactivation. However, there is emerging evidence that macroH2A variants can also be found at sites of open chromatin (Sun et al., 2018; Yildirim et al., 2014). To disambiguate the function of macroH2A2 in epigenetic programs in GBM, we performed the sequencing-based assay for transposase-accessible chromatin (ATAC-seq) in our patient-derived knockdown models. ATAC-seq was performed on two biological replicates for both shScr control cells and *H2A2FY2* knockdown cells. We found that *H2AFY2* knockdown caused both gains and losses of chromatin accessibility (**Figure 4A**), although losses were more frequent and globally resulted in fewer base pairs of DNA in accessible chromatin in knockdown cells (**Figure 4B**). The data support the notion that macroH2A2 contributes to the organization of both accessible and inaccessible chromatin regions, although its contributions to accessible regions are more prominent in our GBM models.

**Figure 4.**
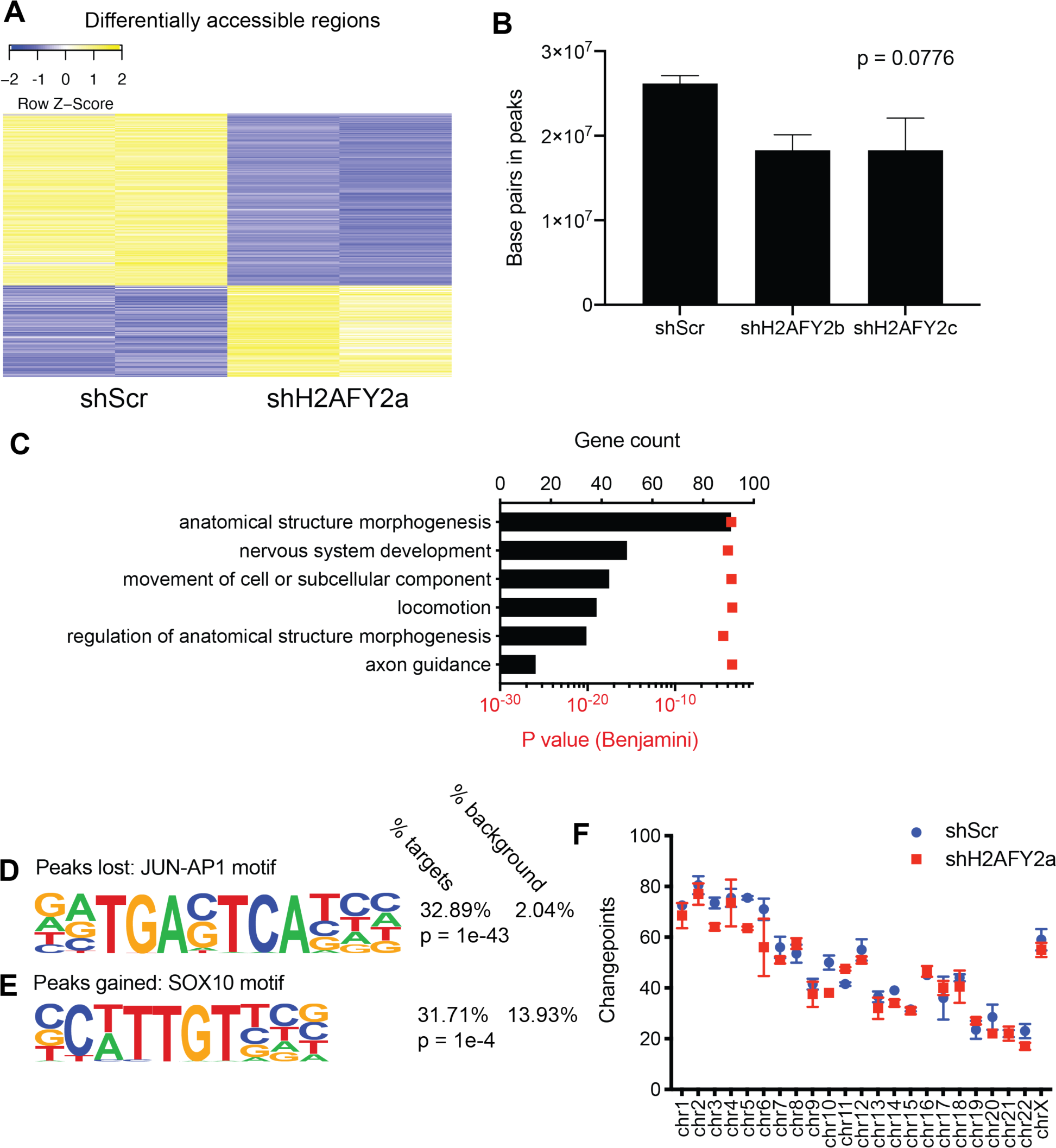
MacroH2A2 contributes to both compacted and accessible chromatin at neurodevelopmental genes in GBM. **(A)** Heatmap of differentially accessible regions (n = 270) in two biological replicates of *H2AFY2*/macroH2A2 knockdown versus control cells. **(B)** Total number of base pairs in peaks of accessible chromatin, as determined by ATAC-seq, in control versus knockdown cells. P value was determined by one-way ANOVA. **(C)** Top process terms resulting from Gene Ontology term analysis of differentially accessible regions. **(D)** Example of a transcription factor motif overrepresented in regions of reduced accessibility in knockdown cells. P value calculated by hypergeometric test. **(E)** Example of a transcription factor motif overrepresented in regions of increased accessibility in knockdown cells. P value calculated by hypergeometric test. **(F)** Changepoint analysis showing number of transitions between closed and accessible chromatin regions in control versus *H2AFY2*/macroH2A2 knockdown GBM cells.

Gene ontology analysis of genes located in differential chromatin accessibility regions upon *H2AFY2* knockdown identified significant enrichment for genes involved in neurodevelopmental pathways, including the terms “nervous system development” and “axonal guidance” (**Figure 4C**). HOMER motif analysis (Heinz et al., 2010) revealed that chromatin accessibility changes occur at sites with significant enrichment for binding sites of transcription factors associated with neurodevelopment. For instance, sites that lost chromatin accessibility had an enrichment of c-JUN motifs (identified at 32.89% of targets compared to 2.04% of background genomic sites, p = 1e-43, hypergeometric test; **Figure 4D**), whereas sites that gained accessibility upon macroH2A2 knockdown had enrichment for SOX10 motifs (identified at 31.71% of targets compared to 13.93% of background genome-wide sites, p = 1e-4; hypergeometric test; **Figure 4E**), a transcription factor associated with neural crest development and oligodendrocyte differentiation (Claus Stolt et al., 2002; Finzsch et al., 2008; Stolt et al., 2004). These findings, together with the developmental regulation of *H2AFY2* we showed above (**Figures 1D,E**), implicate macroH2A2 in fine tuning brain-specific epigenetic and transcriptional programs of self-renewal by modulating chromatin organization.

Next, we investigated whether macroH2A2 levels had a greater impact on *global* chromatin architecture, as we have reported for the histone variant H3.3, compared to its effects on the *local* chromatin environment. We have previously shown that ATAC changepoint analysis provides a measure of global changes in chromatin architecture. We performed changepoint analysis using our ATAC-seq datasets generated with shScr and shH2AFY2 GBM cells and we did not observe large structural differences with the exception of chromosomes 3, 5 and 10 (**Figure 4F**). We conclude that macroH2A2 has dual roles in maintaining compacted and accessible regions without causing large-scale chromatin reorganization. We therefore decided to investigate potential roles of macroH2A2 in shaping the local chromatin environment.

### macroH2A2 represses enhancer function in GSCs

There is very little known on the effects of macroH2A2 on chromatin organization and transcriptional control, and this is particularly true in GBM. We initially hypothesized that macroH2A2 might exert its effects on transcription by modulating chromatin accessibility at gene bodies. Surprisingly, permutation analyses revealed a clear depletion of both gained and lost ATAC peaks in gene bodies and their promoters upon *H2AFY2* knockdown (p = 0.002, **Figure S3**). On the other hand, ATAC peaks that were gained upon knockdown showed only a slight but significant depletion (p = 0.0039) at enhancer elements **(Figure 5A)**, while those lost upon *H2AFY2* knockdown were significantly over-represented at enhancer elements (p = 0.002; **Figure 5B**). Specific peaks gained upon knockdown were identified at putative enhancer regions associated with developmental genes, including *FOXP1* and the oligodendrocyte lineage gene *COL20A1* (**Figure 5C,D)**.

**Figure 5.**
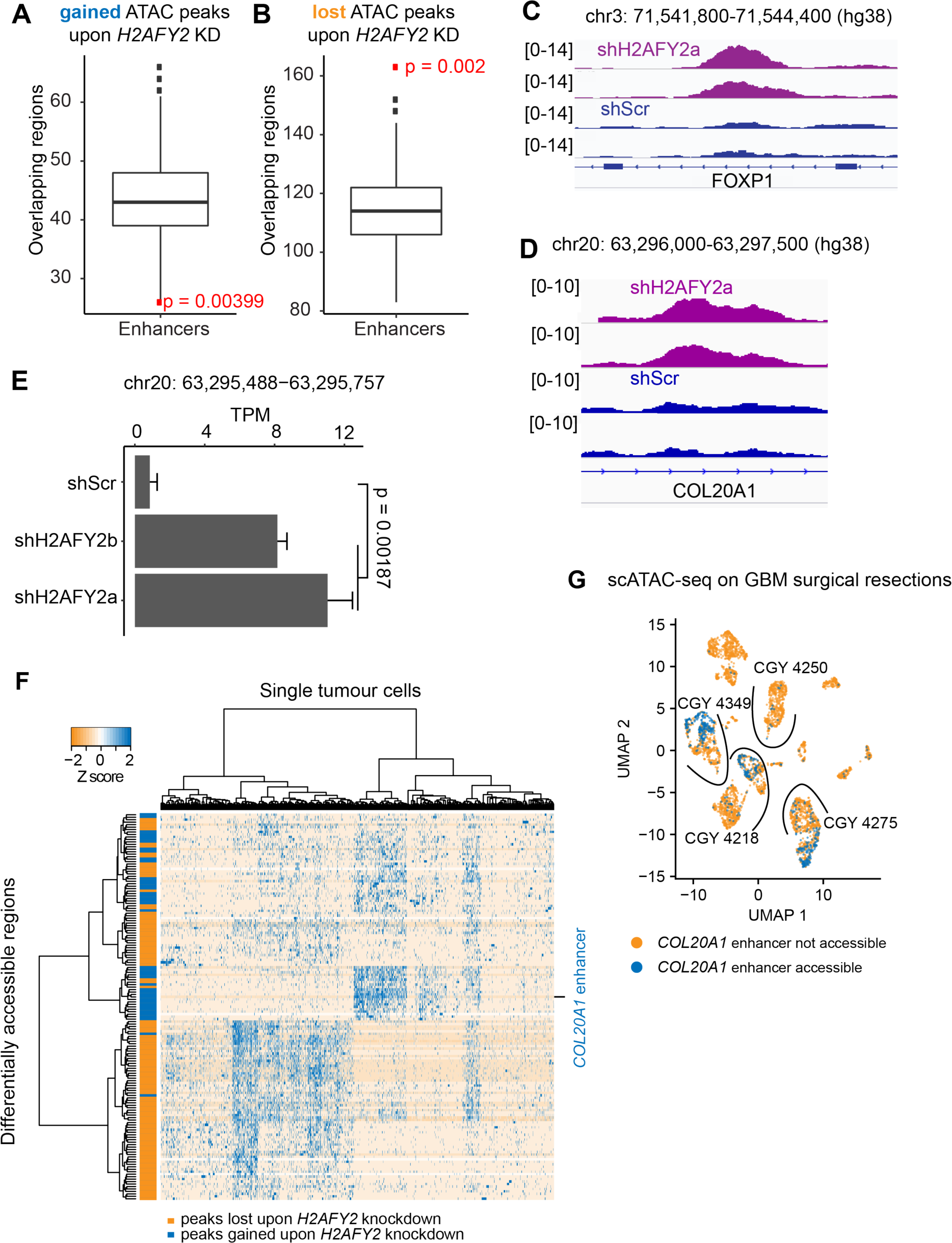
macroH2A2 represses enhancer elements linked to neurodevelopmental genes. (**A-B)** Permutation analysis of accessible regions gained **(A)** and lost **(B)** upon *H2AFY2*/macroH2A2 knockdown at enhancer elements genome-wide. **(C, D)** Examples of ATAC-seq enhancer peaks gained upon knockdown of *H2AFY2*/macroH2A2. **(E)** Expression of eRNA at the *COL20A1* locus shown in **(D)** in control versus knockdown cells. Two biological replicates were used to generate RNA-seq libraries. P value was obtained by two-tailed t-test.. **(F)** Differentially accessible chromatin regions identified upon *H2AFY2*/macroH2A2 knockdown in a scATAC-seq sample from four primary glioblastoma resections. The horizontal axis represents individual cells in the specimens, with differentially accessible regions listed along the Y axis. **(G)** Accessibility at the *COL20A1* enhancer in a scATAC-seq dataset generated from four primary GBM surgical specimens. Samples are separated by arcs.

Active enhancers are transcribed by RNA polymerase II, resulting in the production of enhancer RNA (eRNA) (Li et al., 2016). eRNA levels often closely track the transcription levels of their associated genes (Azofeifa et al., 2018; Hah et al., 2013). We reasoned that if macroH2A2 affects the function of enhancer elements, changes in enhancer activity associated with depletion of macroH2A2 should result in alterations in eRNA transcription levels. Analysis of RNA-seq data from control and *H2AFY2*/macroH2A2 knockdown GBM cells showed significant differential transcription of 33 distinct eRNA transcripts **(Figure S4A-S4E**). Interestingly, we observed a significant transcriptional increase of an eRNA at the *COL20A1* enhancer locus in our knockdown cells **(Figure 5E)**, suggesting that macroH2A2 may play a role in inhibiting this enhancer site. These data are consistent with the effects of *H2AFY2* knockdown on *COL20A1* transcription, and suggest a mechanism by which macroH2A2 represses this gene associated with the oligodendrocytic lineage by compacting the chromatin at its cognate enhancer element.

To further validate the differentially accessible regions we identified in our knockdown cells, we analysed single-cell ATAC-seq (scATAC-seq) we had previously generated on 4 adult GBM primary patient samples (Guilhamon et al., 2021). Adult GBM resections showed accessibility in the majority of differentially accessible peaks gained and lost upon *H2AFY2* knockdown, confirming the relevance of the results generated with our patient-derived models. Gained and lost peaks were accessible in two distinct clusters of cells, suggesting that they define populations characterized by different chromatin states **(Figure 5F)**. The *COL20A1* enhancer was also identified in the primary tumor specimens, and its accessibility was restricted to a single cluster of tumour cells in 2 out of 4 primary samples profiled by scATAC-seq **(Figure 5G; Figure S5A-C)**. All together, our data show that *H2AFY2*/macroH2A2 has an important role in mediating enhancer accessibility in GBM cells, and appears to modulate a regulatory network driving GSC cell identity.

### High-content chemical screen identified compounds that increase macroH2A2 levels

We performed further analyses to explore associations between *H2AFY2* levels and clinical parameters in previously published patient cohorts. Interestingly, we observed that *H2AFY2* levels stratify survival only in patients that received radiation with or without chemotherapy (**Figures 6A**). In all cases, we observed that higher transcription levels of *H2AFY2* were associated with better response to therapy, possibly because of its repressing effects on self-renewal programs, which have been linked to radiation resistance. Because of the positive association between macroH2A2 levels and response to radiation, we reasoned that increasing macroH2A2 levels could lead to “differentiation” of GBM cells and could sensitize tumor cells to standard of care therapy. We therefore set out to perform a chemical screen with the goal of identifying compounds that could increase levels of macroH2A2 **(Figure 6B; Figure S6A**). We screened 182 compounds from the Selleckchem Epigenetic Drug Library using the GE InCell automated confocal microscopy system, coupled with immunofluorescence to assess macroH2A2 levels (**Figure 6C, Figure 6D**). We were able to identify 35 compounds that led to a greater than two-fold increase in the percentage of macroH2A2^+^ GBM cells, including the MLL-menin inhibitor MI-3 and the HDAC inhibitor RGFP966 (**Figure 6E**). We have previously reported that MLL-menin inhibition is a potent antagonist of self-renewal and other stemness-associated properties (Gallo et al., 2015; Lan et al., 2017), and we confirmed here that it has a potent negative effect on cell viability (**Figures 6F**). We validated that MI-3 increases macroH2A2 levels by western blot (**Figure 6G**). Similarly, we found that RGFP966 could reduce cell viability with IC50 < 1 μM (**Figure S6B**), thus supporting the robustness of our screen. We decided to focus our follow up studies on MI-3 because of (1) its potent role at reducing self-renewal in GBM and (2) the translational potential associated with Menin inhibition, as evidenced by the recently-reported development of clinical-grade analogs (Krivtsov et al., 2019) and the active use of the Menin inhibitor KO-539 in a phase 1 clinical trial for patients with refractory or relapsed acute myelogenous leukemia (ClinicalTrials.gov identifier: NCT04067336).

**Figure 6.**
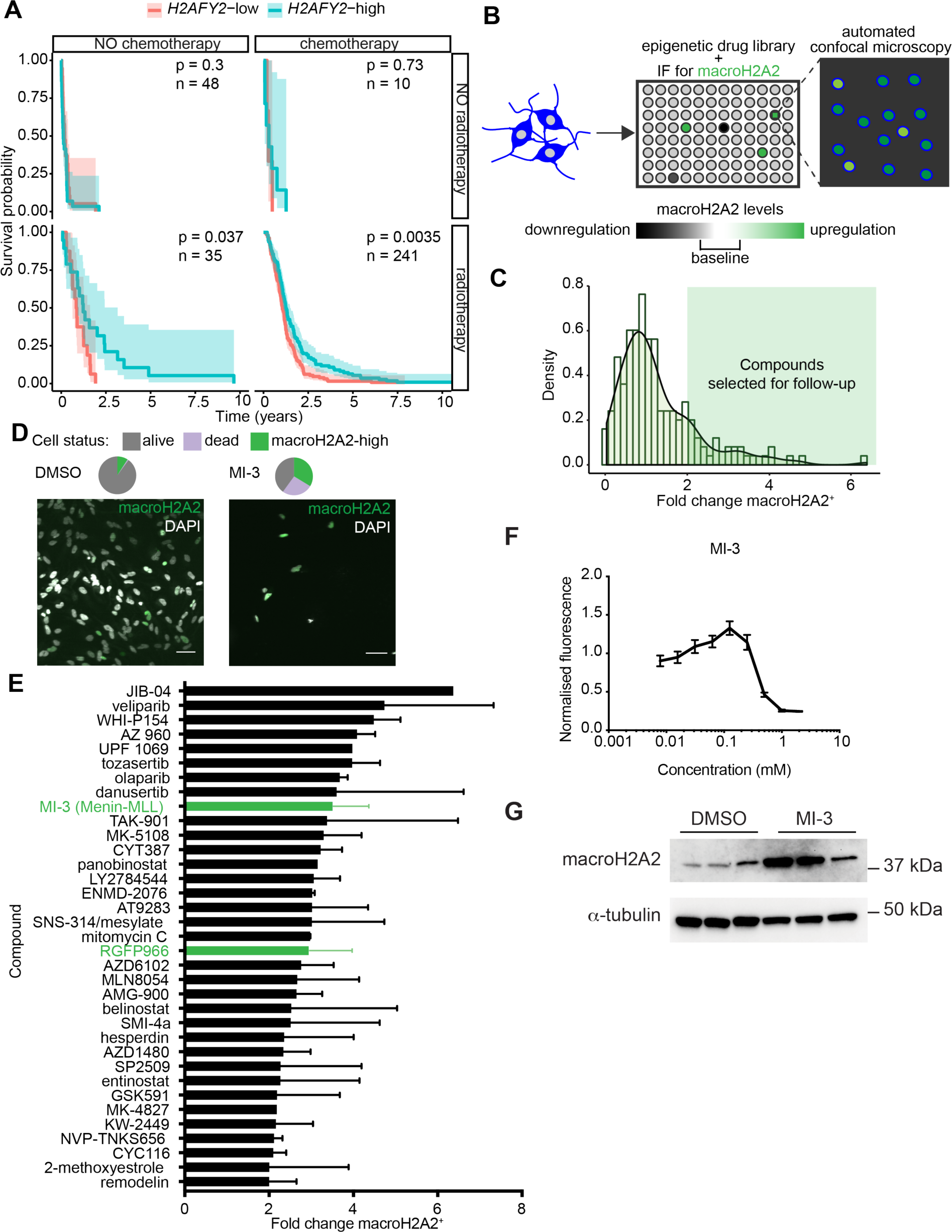
Drug screen to identify epigenetic compounds that elevate macroH2A2 levels in GBM cells. **(A)** Survival analysis of patients in the TCGA GBM cohort segregated by median *H2AFY2* expression and treatment received (none, chemotherapy, radiotherapy, or both). Shaded region represents 95% confidence interval. P value was calculated by log-rank test. **(B)** Diagram summarizing our screening strategy to identify compounds that increase macroH2A2 levels. **(C)** Normalized density of the log fold change of macroH2A2 positive cells for all compounds in the screen. The green shaded region represents compounds with greater than 2-fold change of macroH2A2 positive cells. **(D)** Effects of MI-3 and vehicle control (DMSO) on macroH2A2 protein levels were assessed by immunofluorescence. Scale bars: 50 μm. **(E)** List of all compounds with greater than two fold increase of macroH2A2-positive viable cells upon treatment. **(F)** Dose-response curve for MI-3 using GSC cultures, as measured by the Alamar Blue assay. Experiment performed with six technical replicates per concentration, normalized to DMSO control. Error bars represent standard deviation. The experiment was repeated 2 times. **(G)** Western blot of macroH2A2 levels after 7 days of treatment with 200 nM of MI-3. Three replicates per condition.

### Menin inhibition activates viral mimicry in GBM cells

We then investigated potential mechanisms of action of this epigenetic compound. We treated GBM cells with sub-lethal concentrations (200 nM) of MI-3 or DMSO vehicle control and performed transcriptional studies by RNA-seq (3 biological replicates per condition). Our transcriptomic studies showed that MI-3 treatment results in repression of markers of the oligodendrocytic lineage, including *PDFGRA* and *COL20A1* (**Figure 7A**).

**Figure 7.**
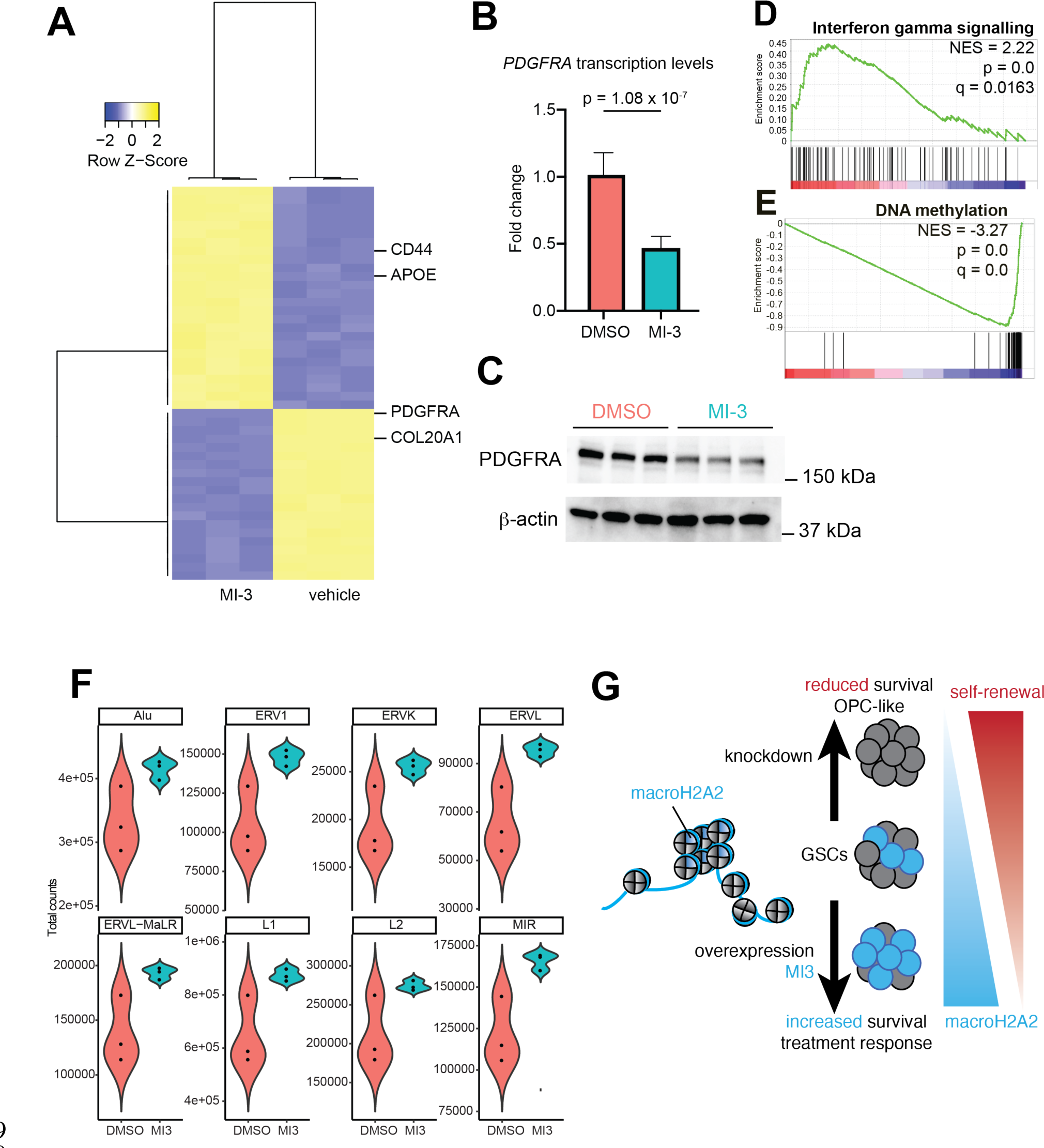
Menin inhibition induces differentiation of GSCs and activates viral mimicry pathways in GBM. (A) Heatmap displaying the top 50 differentially expressed genes between MI-3 and DMSO treated cells based on RNA-seq data (3 biological replicates per treatment). (B) RT-qPCR results showing expression levels of *PDGFRA* in GBM cells after 7 days of MI-3 treatment. Transcription was normalized to actin and GAPDH. P value was calculated by unpaired two-tailed T test. Error bars represent standard deviation. The experiment was repeated two times. (C) Western blot showing levels of PDGFRA and TNFR in MI-3 and DMSO treated GBM cells after 7 days of treatment in vitro. Three biological replicates per condition. The experiment was repeated two times. **(D,E)** GSEA analyses of differentially-expressed genes in MI-3-treated versus vehicle-treated cells. **(F)** Transcriptional levels of repeat elements upon MI-3 treatment were determined by RNA-seq. **(G)** Proposed model for the mechanisms of action of macroH2A2 and Menin inhibition in GSCs.

Repression of *PDGFRA* through MI-3 treatment was confirmed by RT-qPCR (**Figure 7B**) and at the protein level by western blot (**Figure 7C**). Both *PDGFRA* and *COL20A1* were negatively regulated by macroH2A2 (**Figure 3E**), and our data are therefore consistent with MI-3 acting through macroH2A2 to execute chromatin and transcriptional programs, thus validating the premises of our chemical screen.

We performed GSEA and found that differentially-regulated genes following MI-3 treatment were positively associated with signatures of interferon gamma, alpha and beta signaling (**Figures 7D; Figure S7A**) and negatively associated with signatures linked to DNA methylation (**Figure 7E**) and PRC2-mediated methylation of histones (**Figure S7B**). Modulation of interferon signalling related to altered DNA methylation has been reported with epigenetic treatments that elicit viral mimicry in cancer cells (Chiappinelli et al., 2015; Roulois et al., 2015). Viral mimicry is dependent on the transcription of repetitive elements across the genome, particularly ERVL elements (Krug et al., 2019). We therefore re-analyzed our RNA-seq datasets to specifically look at the effects of MI-3 treatment on the transcription of repetitive elements. We found that most differentially transcribed repetitive elements were upregulated by MI-3 treatment, including ERVL family members (**Figure 7F; Figure S7C, S7D**). Our data therefore reveal a role for Menin inhibition in repressing self-renewal programs by upregulating macroH2A2 and potentially activating viral mimicry pathways in GBM **(Figure 7G)**.

## DISCUSSION

Self-renewal is a fundamental property of cancer cells that are therapy-resistant and responsible for long-term tumor propagation. The identification of molecular mechanisms that are responsible for the attainment and maintenance of self-renewal could be key to significantly improve patient outcomes for difficult to treat cancers like GBM (Clarke, 2019). Self-renewal in GBM cells is dependent on the achievement of specific chromatin states and transcriptional profiles through the regulated function of chromatin remodelers, transcription factors and other epigenetic regulators (see for instance Bender et al., 2013; Gallo et al., 2013, 2015; Heddleston et al., 2012; Jin et al., 2017; Miller et al., 2017; Suvà et al., 2009). Interfering with chromatin and epigenetic factors that have key roles in the self-renewal program can destabilize these chromatin states and result in partial or complete loss of self-renewal.

In the present work, we identify macroH2A2 as a key antagonist of self-renewal properties in GBM. This conclusion is supported by multiple orthogonal lines of evidence. First, high *H2AFY2* transcript levels are associated with better response to radiotherapy and longer overall survival in GBM patient cohorts. These effects of *H2AFY2* were recapitulated in our *in vivo* orthotopic patient-derived models, showing a direct effect of this gene in regulating tumor aggressiveness. Second, our *in vitro* and *in vivo* functional assays demonstrated that macroH2A2 has a direct functional role in curbing self-renewal. Our genomic approaches showcased a new function for macroH2A2 in shaping chromatin organization at enhancer elements and regulating the expression of a self-renewal gene network. Overall these results indicate that macroH2A2 has an important effect on the regulation of chromatin and transcriptional dynamics that inhibit self-renewal in GBM.

The development of compounds that target chromatin and epigenetic factors – known as “epidrugs” – has opened new opportunities to pharmaceutically inhibit self-renewal networks in cancer cells. Some of these compounds have been shown to be effective and are currently in use in a subset of patients with certain types of hematological malignancies (Estey, 2013; Fenaux et al., 2009; San-Miguel et al., 2014) We reasoned that if high macroH2A2 levels are associated with better response to radiotherapy, treatment approaches that elevate macroH2A2 levels prior to radiation therapy could improve patient outcomes. We therefore performed a screen to identify compounds that could increase macroH2A2 levels. The Menin inhibitor MI-3 was one of our best hits.

We previously reported that Menin inhibition was a potent repressor of self-renewal in GBM (Lan et al., 2017). Other groups reported effectiveness of Menin inhibition in pediatric glioma models (Funato et al., 2014). However, the epigenetic mechanisms underlying the effectiveness of Menin inhibition in glioma models have not yet been established. Here we show that Menin inhibition acts through at least two complementary but converging mechanisms. First, Menin inhibition results in increased macroH2A2 levels, which in turn repress self-renewal transcriptional networks. Second, Menin inhibition causes activation of transcription of repetitive elements that have been linked to the onset of viral mimicry pathways and lead to the demise of cancer cells (Chiappinelli et al., 2015; Krug et al., 2019; Roulois et al., 2015). Importantly, the Menin inhibitor KO-539 is being tested in a phase 1 clinical trial for patients with refractory or relapsed acute myelogenous leukemia (ClinicalTrials.gov identifier: NCT04067336). A second highly bioavailable Menin inhibitor was recently described and was shown to be effective against patient-derived preclinical models of leukemia (Krivtsov et al., 2019). Targeting Menin is therefore becoming an achievable reality in the clinic, and our findings suggest that this treatment approach should further be tested in preclinical models of GBM.

Because repression of macroH2A2 contributes to aggressive phenotypes in several solid malignancies, it is possible low levels of this histone variant could be used as a biomarker to identify patients that could benefit from more aggressive treatment approaches. These patients could receive clinical-grade Menin inhibitors to increase the levels of macroH2A2, which in turn would repress self-renewal states and curb the aggressiveness of the disease. More studies will be required to explore the translational opportunities presented by Menin inhibition in the context of solid malignancies.

In conclusion, our work characterizes a previously unknown and targetable epigenetic mechanism regulated by the histone variant macroH2A2. Our data underscore the close connection between chromatin and functional states in cancer cells.

## ACKNOWLEDGMENTS

Funding for this work was provided by a Canadian Institutes of Health Research (CIHR) grant (PJT-156278) to MG; a Natural Sciences and Engineering Research Council (NSERC) Discovery grant to MG; a Clinician Investigator Program fellowship to AN; an NSERC studentship to AB. Our research was supported by SU2C Canada Cancer Stem Cell Dream Team Research Funding (SU2C-AACR-DT-19-15) to PBD provided by the Government of Canada through Genome Canada and CIHR, with supplemental support from the Ontario Institute for Cancer Research, through funding provided by the Government of Ontario. SU2C Canada is a Canadian Registered Charity (Reg.# 80550 6730 RR0001). Research funding is administered by the AACR International - Canada, the Scientific Partner of SU2C Canada.

## MATERIALS AND METHODS

### Experimental Model and Subject Details

All specimens and primary cultures generated and used in this study were approved by the Health Research Ethics Board of Alberta and the research ethics board of the Hospital for Sick Children.

GSC primary cultures G523, GSC3 and GSC5 were generated using previously described methods; in brief, primary tumor sample was minced in Accutase (StemCell Technologies), and dissociated with glass beads on a nutator for 30 minutes, followed by centrifugation and resuspension in NS media (Pollard et al. 2009). Cultures were STR genotyped and confirmed to match the patient tissue.

### Cell Culture

Primary glioblastoma cultures were grown in adherent culture on Corning Primaria dishes coated with poly-L-ornithine (Sigma-Aldrich, P4957) and laminin (Sigma-Aldrich, L2020), under standard temperature, oxygen and humidity conditions. Cells were kept in NeuroCult NS-A Basal Medium and Proliferation Supplement (StemCell Technologies, #05751), supplemented with 20 μg/mL rhEGF (Peprotech, AF-100-15), 10 μg/mL bFGF (StemCell Technologies, #78003), and 2 μg/mL heparin (StemCell Technologies, #07980). All cultures were used within the first 20 passages of generation. Adherent cells were disassociated with Accutase (Stemcell Technologies, #07920) and plated onto fresh, coated plates when confluence reached 80/90%. Cell numbers and viability were determined using Countess II (Thermo Fisher Scientific, AMQAX1000).

### Generation of macroH2A2 knockdown cultures

Commercial inducible shRNA constructs (3 for *H2AFY2*, and non-targeting control 1) were obtained from Dharmacon (Horizon Biosciences) and packaged into lentiviral particles. Primary cells were infected with lentiviral particles in the presence of polybrene followed by selection using 1.5 μg/mL puromycin for 72 hours.

### MacroH2A2 overexpression with CRISPRa

#### Generation of an inducible overexpression line

The PB-TRE-dCas9-VPR construct (Addgene #63800) (Chavez et al., 2015) was used to create a stable line using Piggybac transposase (System Biosciences). A total of 500,000 cells were transfected using the using the mouse neural stem cell nucleofection kit (Lonza, VPG-1004) with 0.66 μg of the construct and 0.33 μg of transposase using the Lonza Amaxa Nucleofector I with protocol A-33, followed by selection with hygromycin B at 50 ug/mL for 7 days.

#### Generation of sgRNA plasmids

Guide RNAs were designed using the crispr.mit.edu guide design tool, using as templates the DNAse accessible region directly upstream of the TSS of *H2AFY2*(Kent et al., 2002). The plasmid backbone pLKO-sgRNA-GFP (Addgene #57822) was used and sgRNA plasmids were constructed using BsmBI digestion followed by ligation, as previously described (Heckl et al 2014).

#### Transfection for overexpression experiments

Two million cells containing the dCas9-VPR construct were transfected with a pool of multiple guides targeting *H2AFY2* or a non-targeting control using the Amaxa Nucleofector. Doxycycline (2 μg/mL) was added to media 24 hours after transfection.

### *In vitro* limiting dilution analysis

Cells were plated on uncoated low-adhesion 96-well plates in a two-fold dilution series spanning from 2000 down to 4 cells per well in NeuroCult NS-A media (StemCell Technologies, #05751) containing doxycycline at 2 μg/mL, with 6 replicates per concentration. Sphere formation frequency was estimated using ELDA (Hu and Smyth 2009). Sphere formation was scored on day 7 and day 14.

### RT-qPCR

RNA samples were used to generate cDNA using the SuperScript II kit (Invitrogen) and poly-A primers. PCR was performed using the SSOFast EvaGreen Supermix (BioRad # 1725201) on the BioRad CFX with all samples in triplicate. Results were analysed using the delta Ct method.

### Western blot

Protein concentration of samples was determined using the DC (detergent compatible) protein assay (Bio-Rad, #5000112). Samples were prepared in a total volume of 20 μL at 15 μg/μL in Laemmli loading buffer. Samples were run on 7.5% Mini-PROTEAN gels for cytoplasmic proteins and 12.5% Mini-PROTEAN gels for histones (Bio-Rad, #4568025). Primary antibodies used: Rabbit anti-mH2A2 (Invitrogen; PA5-57437), mouse monoclonal anti-β-ACTIN (Sigma-Aldrich, A5441, Lot# 127M4866V, Clone AC-15) at 1:1000), rabbit anti-PDGFR*α* (CST #3164), rabbit monoclonal TNF-R1 (CST #3736). Secondary antibodies used: Goat anti-rabbit IgG H&L (HRP) (Abcam, #6721, Lot# GR3192725-6) 1:20,000, Goat anti-mouse IgG H&L (HRP) (Abcam, #6789).

### Mouse intracranial orthotopic xenografts

#### Mouse survival

For each mouse, 100,000 tumor cells (control or knockdown) in PBS were stereotactically injected into the forebrain (Location: 2.0-3.0 mm to the right of bregma, 1.0 mm anterior to coronal suture) of 3 month old female NSG mice (Jackson Stock no 005557), using a 30 gauge needle. Mice were fed 2 mg/mL doxycycline in 2% sucrose water solution. Endpoint was reached once mice showed signs of disease, including ataxia, hunching, domed heads, kyphosis, paresis and lethargy.

#### Orthotopic limiting dilution assay

Stereotactic injections of 100,000, 10,000 or 1,000 cells were performed into the right forebrain of NSG mice as described above. Mice were sacrificed at endpoint as in the previous experiment.

### ATAC-seq

#### Experimental method and sequencing

ATAC seq was performed using the Omni-ATAC protocol (Corces et al., 2017). In brief, 50,000 cells were harvested fresh from culture from two biological replicates, lysed on ice, spun at 4 C, and treated with Tn5 transposase (Illumina) for 30 minutes at 37 C. Libraries were amplified using the NebNext HiFi polymerase mastermix (New England Biolabs) and standard Illumina Nextera primers, and DNA was purified using the MinElute kit (Qiagen). Sequencing was performed on a NextSeq 500 using 150 cycles of paired end 75 bp sequencing at the Centre for Health Genomics and Informatics (University of Calgary).

#### ATAC-seq data analysis

ATAC-seq data was aligned to hg38 using bwa mem (Li and Durbin, 2009). Following this, samtools was used to remove the mitochondrial and Y chromosome reads, and reads mapping to genome blacklists (Li et al., 2009). Duplicates were removed using picard. Peak assignments were generated using macs2 (Zhang et al., 2008). Differential peak calls were performed by generating a pileup of counts of all consensus peaks from all samples, transforming the counts to counts per million using edgeR, and running these normalized counts through DESeq2 (Love et al., 2014). Peaks with a fold change of > 1.5 were considered significantly altered. Changepoint analysis was run on a pileup of ATAC signal reads binned into 1 mb bins using bedtools, using the R package *changepoint*, as previously described (Gallo et al., 2015). Permutation analysis was performed using the R package regioneR (Gel et al., 2016). Gene ontology analysis was performed using DAVID 6.8 (Huang et al., 2009b, 2009a).

### RNA extraction and RNA sequencing

#### Sample preparation and sequencing

For macroH2A2 knockdown RNA-seq, stably transduced cells were induced with 2 μg/mL doxycycline and grown for 7 days in culture and processed in biological duplicates. For MI-3 vs DMSO treated cells, G523 cells were treated with either 200 nM MI-3 or DMSO for 7 days and processed in biological triplicate. Cells were harvested using Accutase, and RNA was extracted using the RNEasy Mini kit (Qiagen) as per manufacturer instructions. Libraries were constructed at the Center for Health Genomics and Informatics using the NEBNext Ultra II Directional RNA Library prep kit (New England Biolabs) with ribosomal RNA depletion. Samples were sequenced on a NextSeq 500 for 150 cycles in single-end mode for mH2A2 knockdown, and paired-end mode for MI-3.

#### RNA-seq analysis

Samples were pseudoaligned to the human transcriptome (GRCh38.rel79) using kallisto, and differential analysis was performed using sleuth (Bray et al., 2016; Pimentel et al., 2017). GSEA was performed using a ranked list of all genes generated using sleuth. For analysis of eRNAs, the Fantom5 CAGE (Andersson et al., 2014) consensus list of enhancers, as well as a custom list of predicted enhancers based on G523 ATAC-seq and H3K27ac consensus peaks, was used to construct a custom pseudotranscriptome, which was analysed using kallisto and sleuth in a similar fashion. Analysis of repeat expression was performed using REdiscoverTE (Kong et al., 2019). Heatmaps were constructed with R using the heatmap.2 function from the package gplots.

### Dose-response curves

Cells were plated at 4000 cells/well in NS media into laminin-poly-L-ornithine coated 96-well plates (Corning). The compounds MI-3, RGFP-966 and AZD6102 (Selleckchem) were tested at different concentrations (range: 2 μM to 8 nM in serial dilutions) with six technical replicates per dose, and a DMSO control. Cell viability was assessed on day 7. Alamar blue (Thermo Fisher Scientific, Cat# DAL1025) was added and cells were incubated at 37°C in the dark for 4 hours. Fluorescence was measured on the Spectramax spectrophotometer and was normalized to the DMSO control.

### Immunofluorescence High-Content Drug Screening

#### Screen procedure

G523 cells were plated at a density of 5,000 cells per well in a 96-well optical plate coated with PLO-laminin as per protocol. An epigenetic drug library (Z195677-L1900; Selleckchem) was added at 1 μM to plates in triplicate, with DMSO used as a control. Cells were incubated for 10 days and fixed in 4% PFA for 10 minutes followed by storage at 4°C until imaging.

#### Immunocytochemistry and imaging

Plates were blocked in 5% BSA in PBS with 0.1% Triton- X100 for 1 hour at room temperature. Staining was performed overnight at 4°C in the same media with the macroH2A2 antibody (Invitrogen; PA5-57437) at a dilution of 1:250, and a fluorescently conjugated secondary antibody (Invitrogen Goat anti-rabbit IgG A568), and a DAPI counterstain (1:1000; Thermo Fisher #62248). Washes were performed using PBS with 0.1% Triton-X100. Samples were left in the plates in PBS and imaged using the GE InCell 6000 with a 60x objective in 3 different planes of section.

#### Image analysis

Images for each well were stitched together using Fiji and the grid/collection stitching plugin (Preibisch et al., 2009). The DAPI channel was used to generate a list of nuclei and Fiji was used to measure intensity for both DAPI and the GFP channel using an automated custom script. Data was then further analysed using R, where data for all compounds on one plate was pooled together, and k-means clustering using brightness and shape parameters was used to stratify live/dead cells and mH2A2 positive or negative cells. These categories were then used to separate cells in each well as alive and mH2A2-negative, alive and mH2A2-positive, and dead for subsequent analysis. Percentages of mH2A2-positive cells were then compared to DMSO control to calculate a fold change.

## Data availability

All RNA-seq and ATAC-seq data will be deposited to the Gene Expression Omnibus (GEO) upon publication of the manuscript. The scATAC-seq data of adult GBM has been published and is available in GEO (accession: GSE139136) (Guilhamon et al., 2021). Analysis code is available upon request.

## Supplementary Figures

**Supplementary Figure S1, related to Figure 1.**
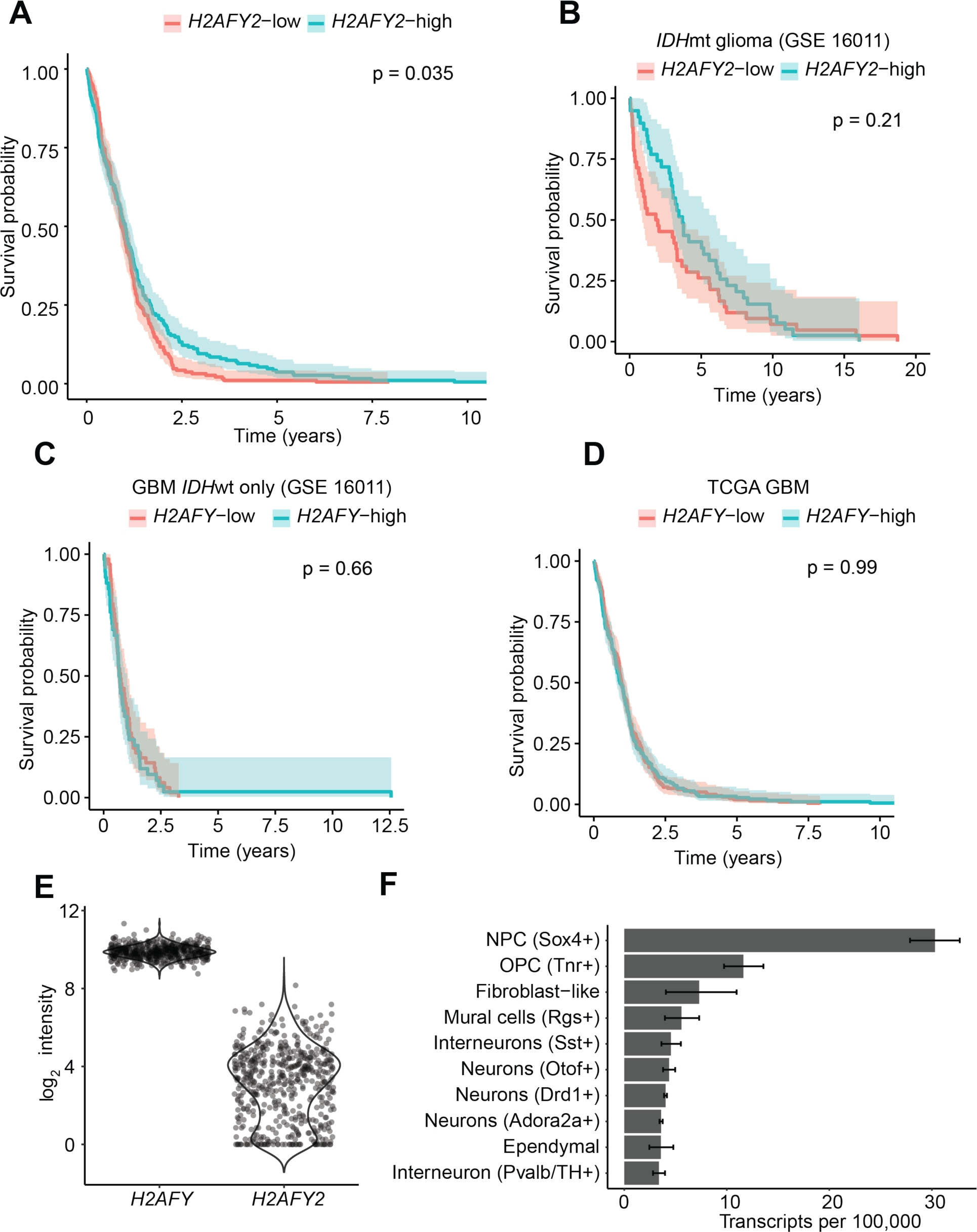
**(A)** Overall survival of the TCGA GBM cohort stratified by median *H2AFY2* level. P value calculated by log-rank test. Shaded region represents 95% confidence interval. **(B)** Overall survival for *IDH* mutant gliomas in GSE 16011 stratified by median *H2AFY2*. P value calculated by log-rank test. Shaded region represents 95% confidence interval. **(C)** and **(D)** Overall survival of *IDH* wildtype GBM (GSE 16011) and TCGA GBM stratified by median *H2AFY* level. P value calculated by log-rank test. Shaded region represents 95% confidence interval. **(E)** Expression data for *H2AFY* and *H2AFY2* in the TCGA GBM cohort. **(F)** Single cell expression data for *H2afy2* in mouse striatum (DropViz mouse datasets).

**Supplementary Figure 2, related to Figure 1.**
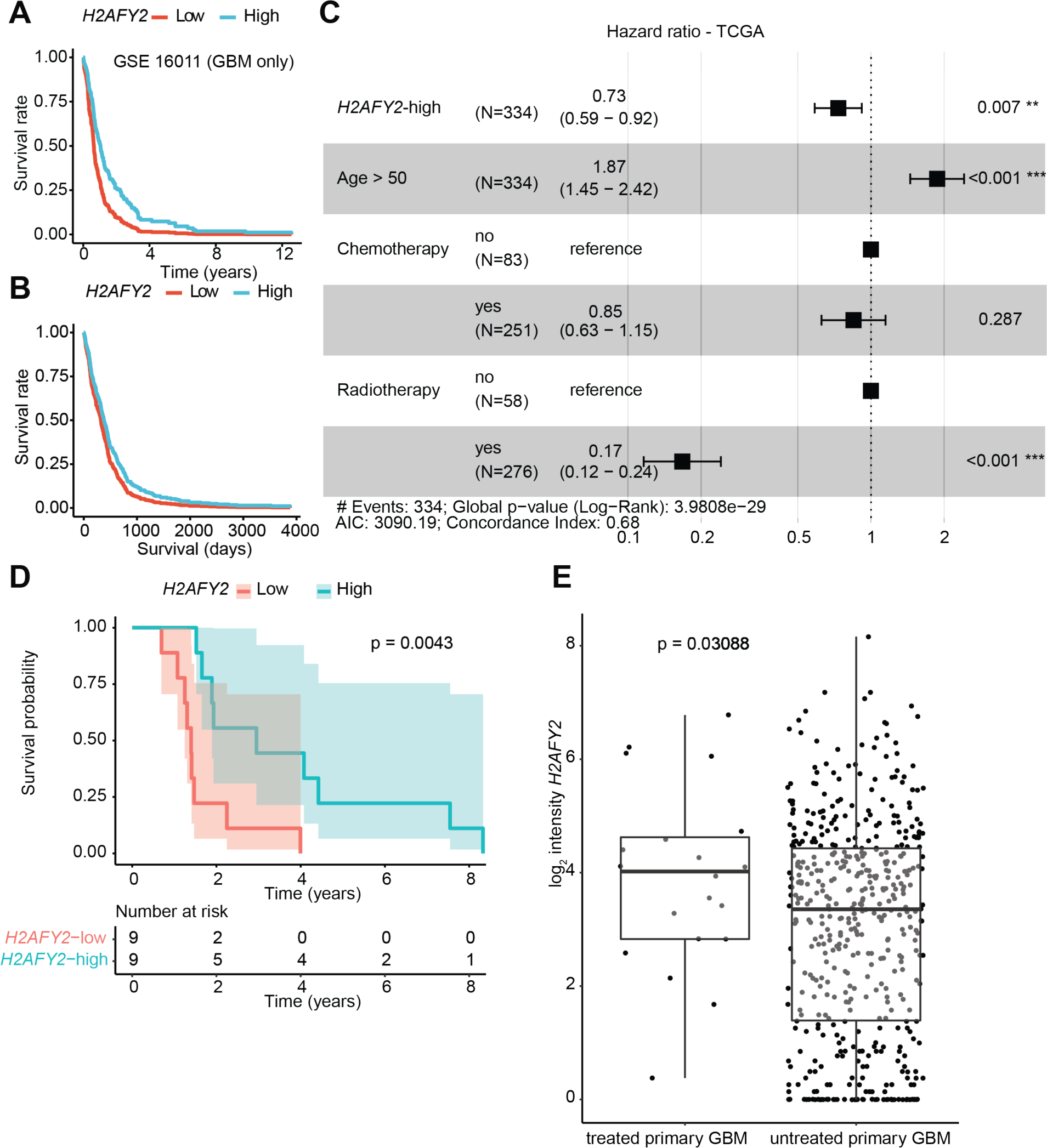
Multivariate cox regression of *H2AFY2* expression in GSE 16011 **(A)** and TCGA GBM **(B)**. **(C)** Cox multivariate regression and hazard ratio for *H2AFY2* expression status with age and chemotherapy and radiotherapy status in TCGA GBM. **(D)** Kaplan-Meier survival status for patients with recurrent glioblastoma in TCGA cohort (shaded region represents 95% confidence interval; p value calculated by log-rank test). **(E)** Comparison of expression levels for *H2AFY2* in TCGA data for untreated versus treated primary glioblastoma (p value: two-tailed T test with Welch’s correction; whiskers represent 95% confidence interval).

**Supplementary Figure S2, related to Figure 2.**
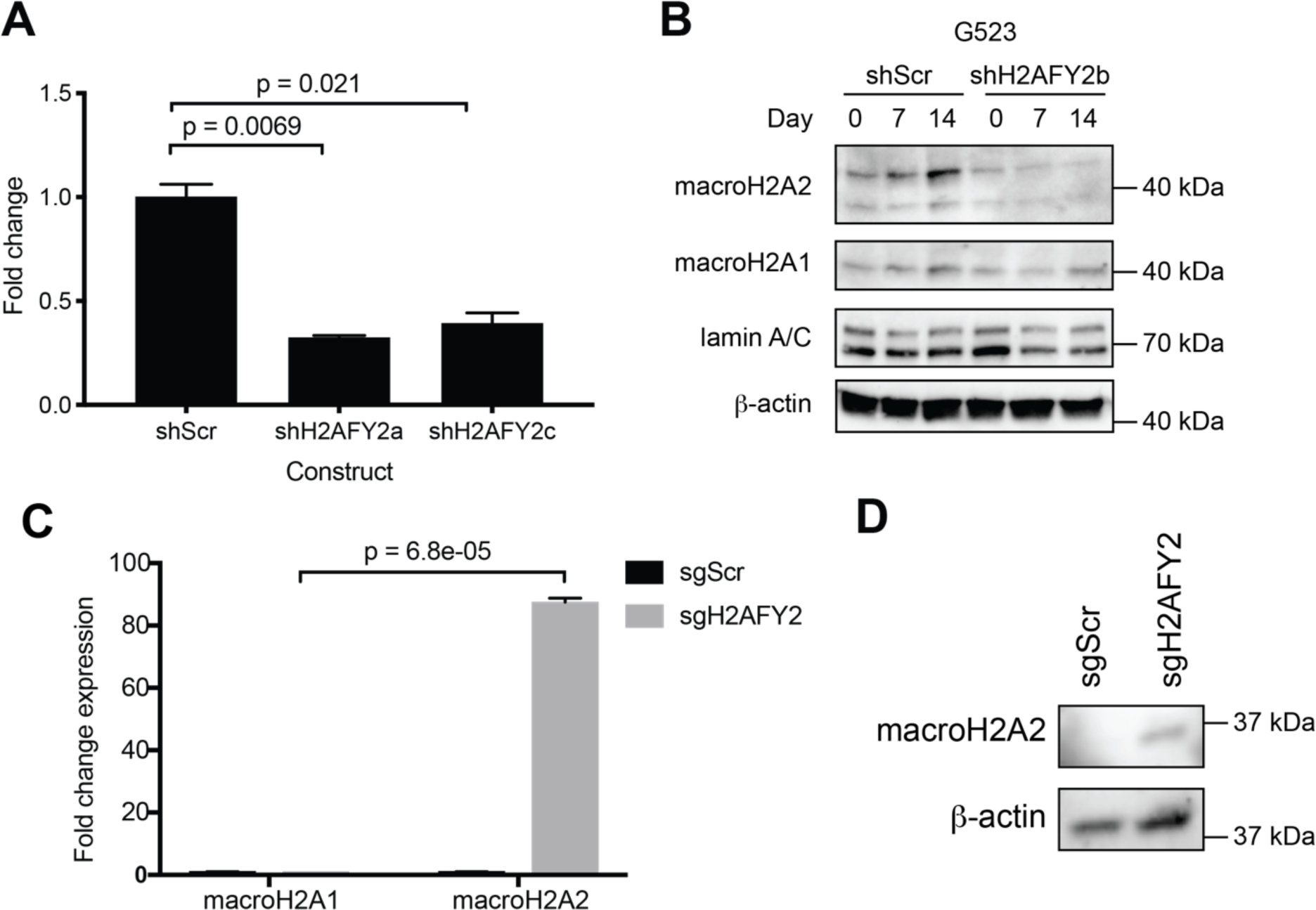
**(A)** RT-qPCR of macroH2A2 expression after 48 hours of doxycycline induction for two distinct hairpins. Expression normalized to actin and GAPDH. P value calculated by unpaired two-tailed T test. Error bars represent standard deviation. Experiment repeated three times. **(B)** Western blot of macroH2A2 and macroH2A1 after 7 or 14 days of activation with doxycycline. Experiment repeated two times. **(C)** RT-qPCR of macroH2A1 and macroH2A2 CRISPR activation in a second primary GSC culture with Expression normalized to actin and GAPDH. P value calculated by unpaired two-tailed T test. Error bars represent standard deviation. Experiment repeated two times. **(D)** Western blot of macroH2A2 levels after 7 days of CRISPR activation in a macroH2A2-low GSC culture. Experiment repeated two times.

**Supplementary Figure S3, related to Figure 5.**
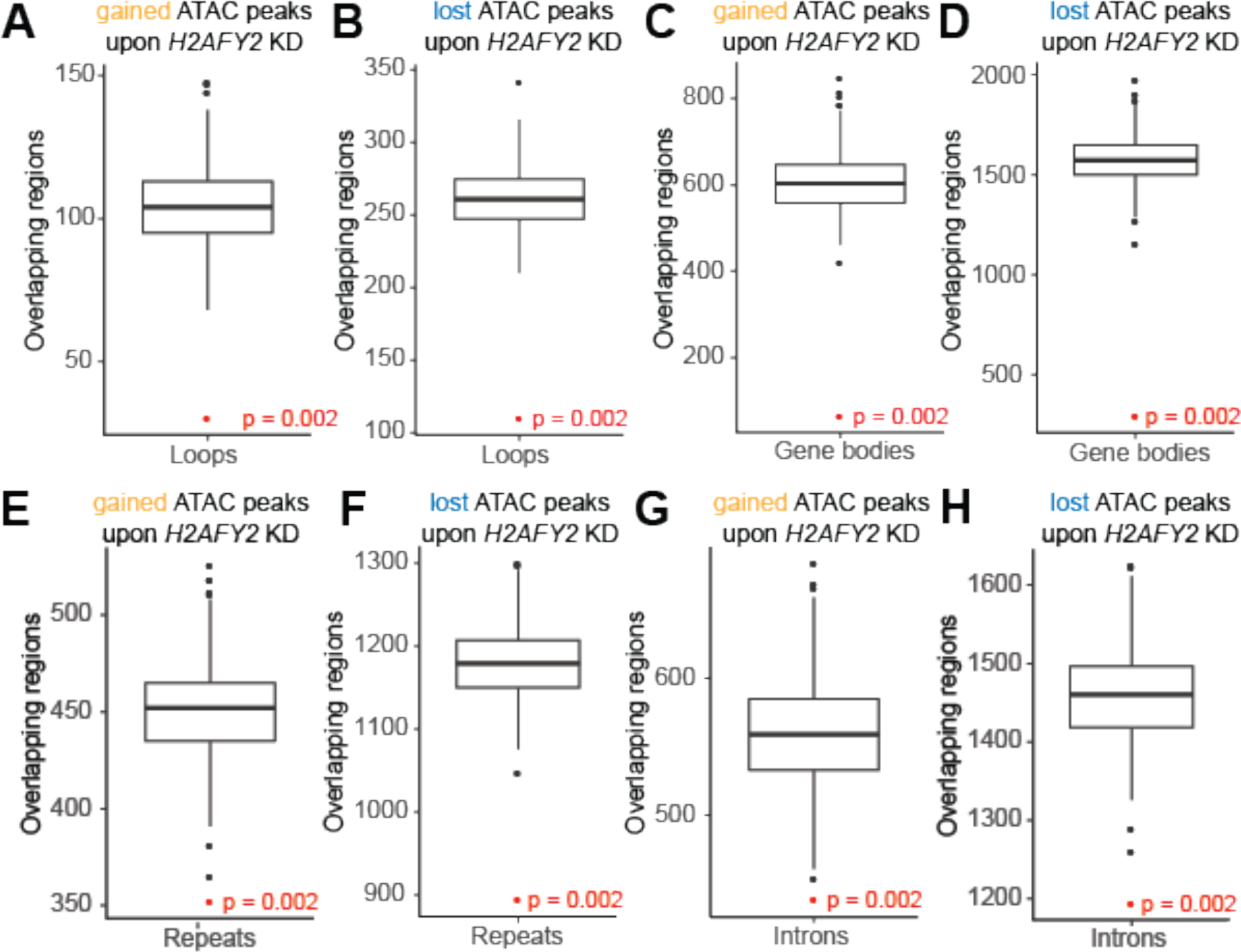
Permutation analysis of regions with at least 1.5 fold log2 change in accessibility upon *H2AFY2* knockdown with **(A-B)** DNA loop boundaries (Johnston et al 2019); **(C-D)** Gene bodies; **(E-F)** repeat regions from the RepeatMasker database; **(G-H)** Introns. Regions of accessibility gain and loss were analysed separately. P value from 500 permutations.

**Supplementary Figure S4, related to Figure 5.**
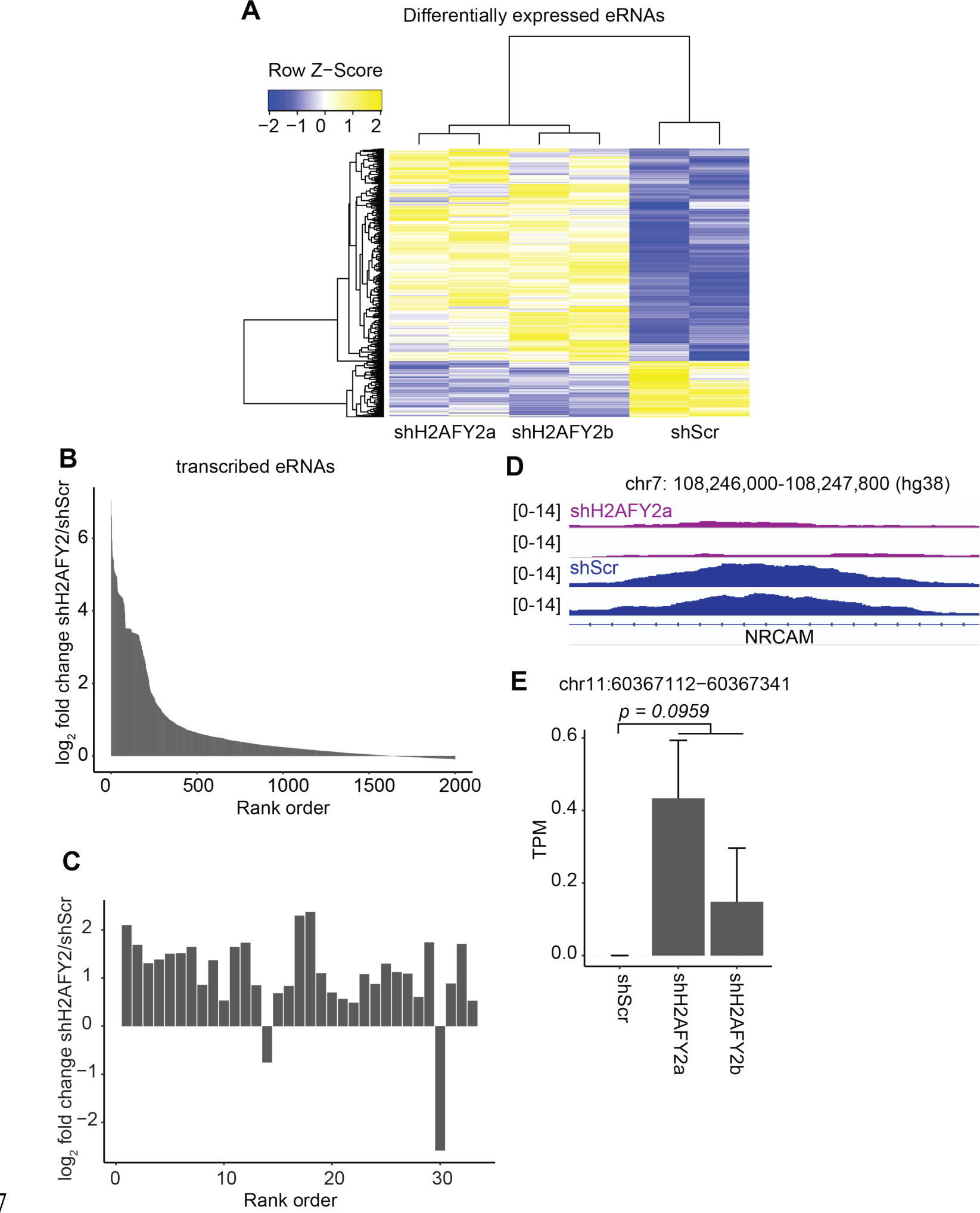
**(A)** Differential expression of top 200 differentially expressed GBM-specific putative eRNAs between *H2AFY2* knockdown and control cells. **(B)** Top 2000 differentially transcribed GBM-specific eRNAs between *H2AFY2*/macroH2A2 knockdown and control cells, ranked by fold change. **(C)** Close up of significantly differentially transcribed eRNA, ranked by p value. **(D)** Example of a representative ATAC-seq enhancer peak lost upon *H2AFY2*/macroH2A2 knockdown. **(E)** Example of eRNA at an enhancer locus with increased accessibility. P value calculated by unpaired T test. Error bars represent standard deviation.

**Supplementary Figure S5, related to Figure 5.**
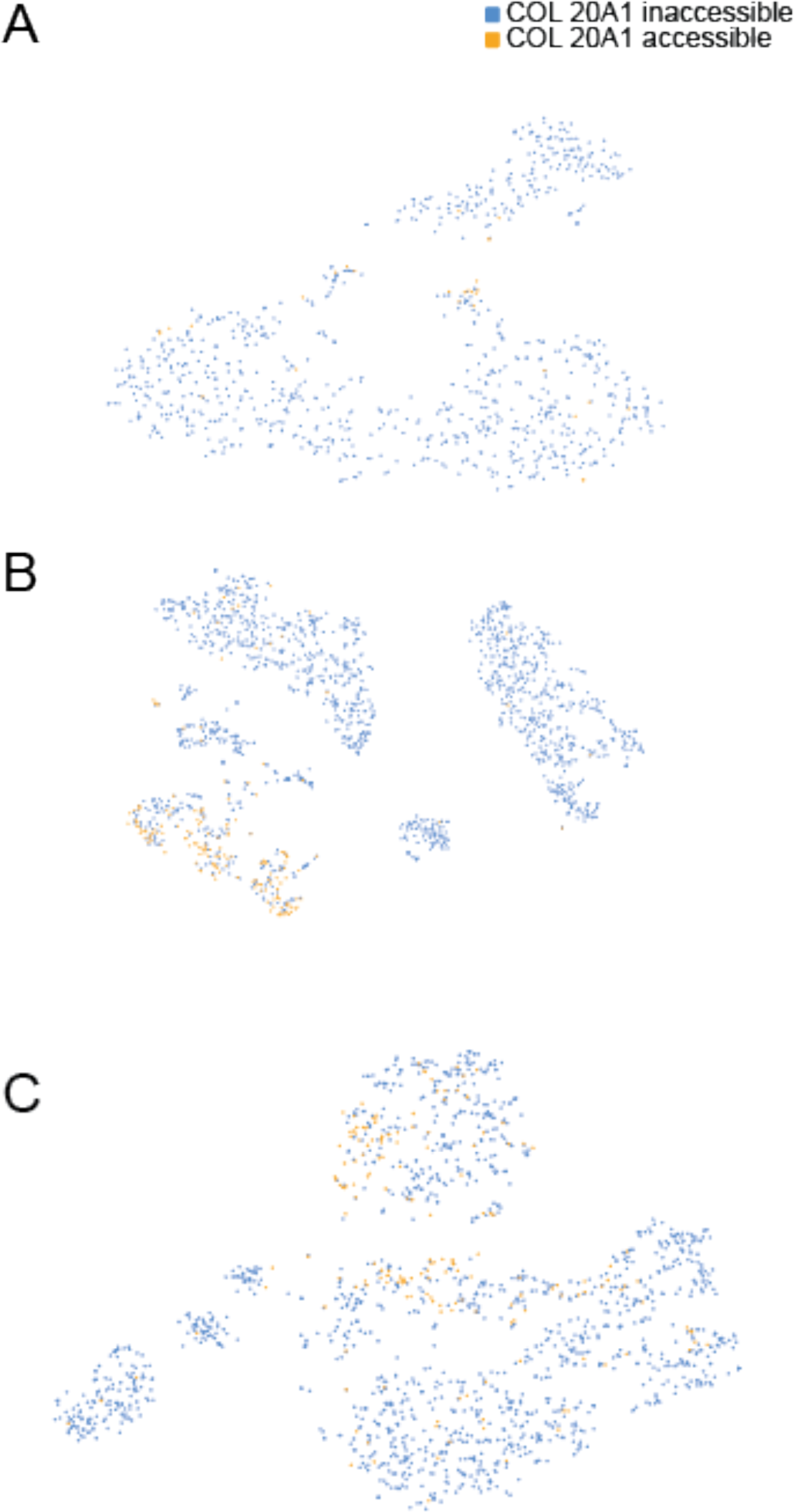
**(A-C)** tSNE plots of scATAC-seq data from three different primary GBM resectionsAccessibility at the *COL20A1* enhancer locus is highlighted in orange.

**Supplementary Figure S6, related to Figure 6.**
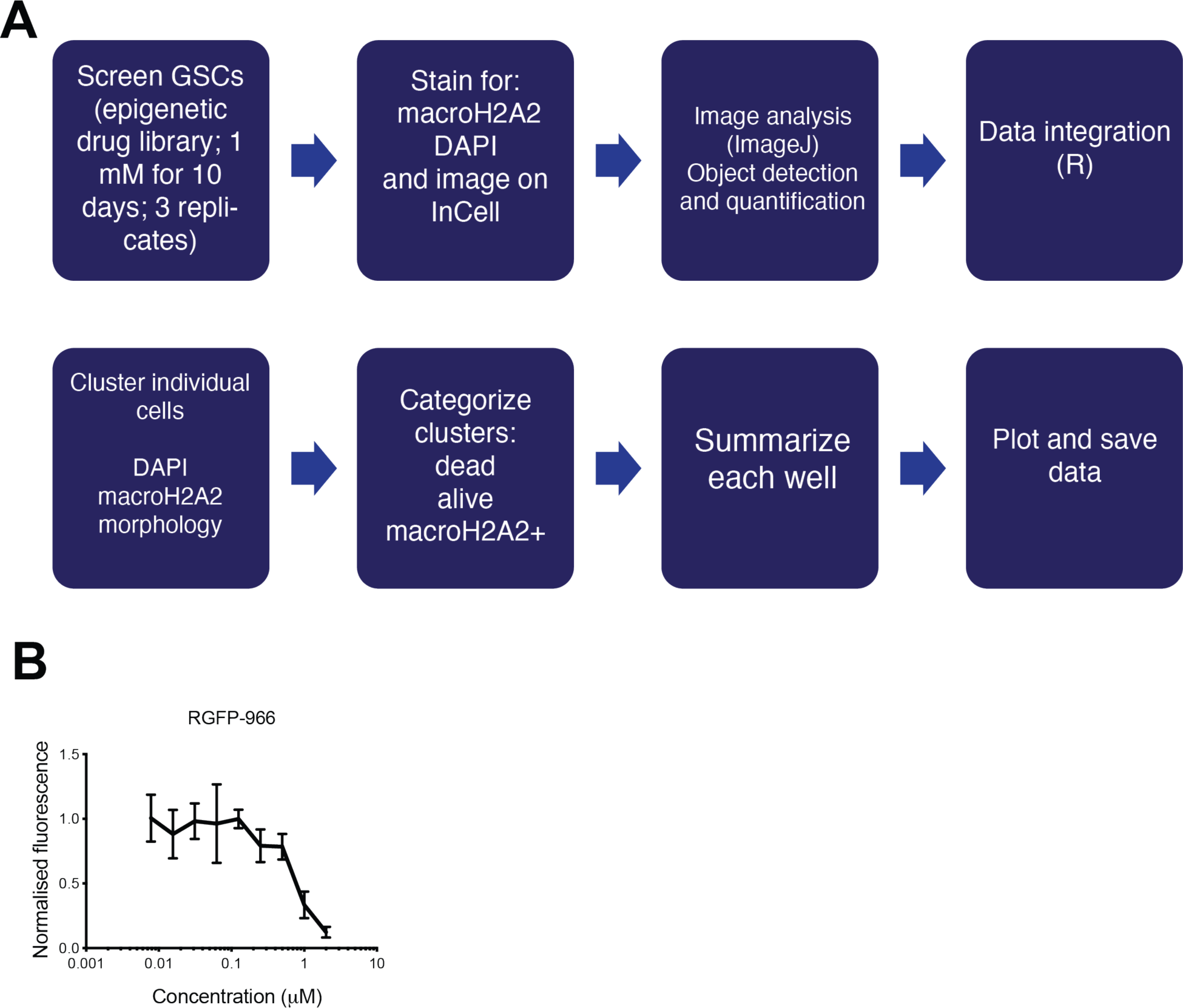
**(A)** Overview of high-content screening strategy. Alamar blue *in vitro* dose-response curve for RGFP-966 in G523 cells. Six technical replicates per concentration. Error bars represent standard deviation. Experiment repeated two times.

**Supplementary Figure S7, related to Figure 7.**
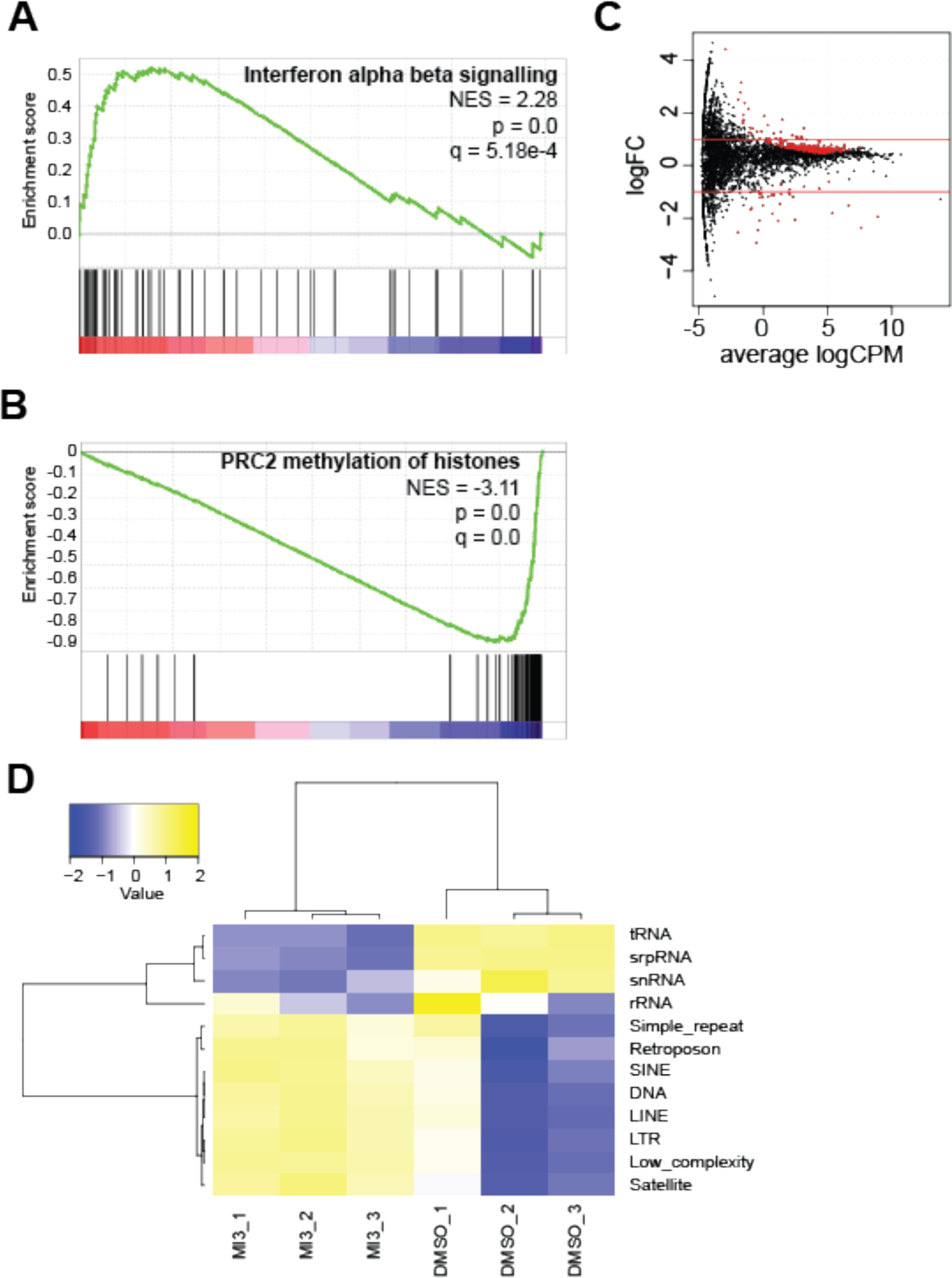
**(A, B)** GSEA results showing altered interferon signalling and methylation signatures upon MI-3 treatment compared to DMSO control. **(C)** Volcano plot showing expression of repeat elements in MI-3 treated cells versus control; points in red represent statistically significant changes (q value < 0.01). Three biological replicates per condition. **(D)** Heatmap showing differential expression of repeat families between vehicle and MI-3 treated cells. Three biological replicates per condition.

